# The reticulocyte restriction: invasion ligand RBP1a of *Plasmodium vivax* targets human TfR1, prohibitin-2, and basigin

**DOI:** 10.1101/2025.05.01.651667

**Authors:** Jessica Molina-Franky, Daniel Röth, Monica Ararat-Sarria, Manuel Alfonso Patarroyo, Markus Kalkum

## Abstract

*Plasmodium vivax* is the most widespread malaria species with predominance in Latin America and Southeast Asia. Devising effective controls against *P. vivax* infections presents unique challenges due to the parasite’s exclusive invasion of reticulocytes, a transient developmental stage of red blood cells. The ephemeral nature and limited availability of reticulocytes complicate efforts to establish continuous *in vitro* cultures to study the parasite’s biology and receptor-ligand interactions. Here, the potential of the erythroid cell lines JK-1 and BEL-A as alternatives to reticulocytes was confirmed through successful parasite invasion, marking the first report of *P. vivax* entry to BEL-A cells. Furthermore, LC-MS was used to quantitatively compare the membrane proteomes of these cell lines with those of reticulocytes and mature erythrocytes. This analysis revealed significant similarities between the membrane proteomes of JK-1, BEL-A, and reticulocytes. In addition to the known *Plasmodium* receptors, transferrin receptor protein 1 (TfR1 or CD71), CD98hc, and basigin (BSG), potential receptor candidates involved in the parasite’s invasion pathway were identified, including prohibitin-2 (PHB2), CAT-1 (SLC7A1), ATB(0) (SLC1A5), CD36, integrin beta-1 (ITGB1), and Metal transporter CNNM3. Proximity labeling using TurboID enabled the identification of specific interactions between PvRBP1a_158-650_ and TfR1, BSG, and prohibitin-2 in the erythroid cell lines. Notably, this is the first report of prohibitin-2 as a receptor for *P. vivax*. These findings advance the understanding of *P. vivax* receptor-ligand interactions and underscore the potential of JK-1 and BEL-A cell lines as alternative models for the study of parasite biology.

**Author Summary:** *Plasmodium vivax* is a major cause of malaria, particularly in Latin America and Southeast Asia. Unlike other malaria parasites, it infects only immature red blood cells (reticulocytes), which are rare and short-lived, making laboratory studies challenging. No continuous *in vitro* culture system currently exists for *P. vivax*, limiting research into its invasion mechanisms. In this study, we evaluated two erythroid cell lines, JK-1 and BEL-A, as potential reticulocyte surrogates. We demonstrate that *P. vivax* can invade both cell types, including BEL-A cells, which had not previously been tested. Comparative analysis revealed key differences in surface protein expression between these cell lines and reticulocytes, compared to mature red blood cells, identifying seven potential parasite-receptor proteins. We further applied a proximity-based protein tagging approach to identify host-parasite interactions mediated by a specific *P. vivax* protein ligand, RBP1a. This technique revealed a potential role for prohibitin-2, TfR1, and basigin in mediating *P. vivax* entry into red blood cells by RBP1a binding. The role of prohibitin-2 as a Plasmodium receptor is a novel Discovery. Our results highlight JK-1 and BEL-A cells as valuable tools for *P. vivax* research and suggest new avenues for investigating host-pathogen interactions, with implications for malaria treatment and vaccine development.

## Introduction

Malaria remains a major global health issue, with 263 million cases reported in 2023. *Plasmodium vivax*, one of the six human-infecting *Plasmodium* species, is noteworthy for its global distribution. In the Americas, most malaria cases are due to *P. vivax* (72.1% in 2023) [1]. Its blood stage form, the merozoites, interact with red blood cell (RBC) receptors leading to RBC invasion [2–4]. However, *P. vivax* exclusively infects reticulocytes [5], immature, short-lived precursors of erythrocytes, posing a significant challenge to research progress. In contrast, *P. falciparum* has been extensively researched due to the availability of a well-established *in vitro* culture system for over 40 years [6]. This critical disparity highlights the urgent need to develop alternative research models for *P. vivax*.

The mechanism of *P. vivax* reticulocyte invasion remains unclear. It was previously believed that *P. vivax* exclusively targeted reticulocytes through the Duffy antigen receptor for chemokines (DARC) and Duffy binding protein (PvDBP) interaction [7], as individuals with the Fy(a-b-) mutation in West Africa were resistant to the infection. However, DARC is present on both reticulocytes and erythrocytes, and *P. vivax* infections have been documented in Duffy-negative populations [8–10]. Current evidence suggests that the reticulocyte-binding protein (RBP) family may play a role, interacting with transferrin receptor 1 (TfR1) and CD98 heavy chain (SLC3A2), which are lost during the maturation of reticulocytes to erythrocytes [2,3,11]. This implies that *P. vivax (*Pv)RBP family proteins may specifically target receptors unique to the reticulocyte membrane. Our previous studies on PvRBP1 of the *P. vivax* strain Belem (GenBank AAA29743.3) identified eleven high-affinity reticulocyte binding peptides (HABPs) corresponding to residues 158–653 of PvRBP1a in the *P. vivax* Salvador I strain (GenBank AAS85749.1). Among these, HABP 3742 (KLLGEEISEVSHLYV) and HABP 3459 (KEILDKMAKKVHYLK) exhibited dissociation constants (*K*d) of 131 nM and 155 nM, respectively [12]. Additionally, an extracellular portion of PvRBP1a, residues 157-650, binds strongly (∼50%) to reticulocytes and moderately (∼20%) to erythrocytes [13]. The identity of PvRBP1a_157-650_ binding-receptors within the reticulocyte membrane has been unclear. Therefore, this study evaluated the erythroleukemic cell line JK-1 [14] and the <u>B</u>ristol <u>E</u>rythroid <u>L</u>ine <u>A</u>dult (BEL-A) [15] as surrogates for reticulocytes, examining their susceptibility to *P. vivax* invasion. A comparison of the cell lines’ membrane proteomes revealed similarities with those of reticulocytes, and dissimilarities with erythrocyte membrane proteomes, thereby identifying potential *P. vivax* receptors. Furthermore, TurboID proximity labelling implied specific interactions of PvRBP1a_158-650_ with prohibitin-2 (PHB2), TfR1, and basigin (BSG). These interactions were confirmed by ELISA, highlighting key molecular determinants of *P. vivax*’s reticulocyte tropism.

## Results

### *P. vivax* can invade the erythroid cell lines BEL-A and JK-1

Cultured BEL-A and JK-1 cells were successfully invaded by *P. vivax* from a validated mono-infected malaria patient’s blood sample. The experiment required forgoing enrichment of the RBCs to 4.0% parasitemia (Fig 1 A-E). Parasite invasion was confirmed by immunofluorescence microscopy, detecting the intracellular presence of *P. vivax* lactate dehydrogenase (PvLDH), which all blood stages of the parasite are known to express [16]. PvLDH was detected in the positive control of infected reticulocytes, and in BEL-A and JK-1 cells, incubated with infected reticulocytes (Fig 2, FITC). The PvLDH signal was absent from non-infected control cells. Moreover, the presence of hemozoin (Hz) pigment, characteristic for hemoglobin consumption by *Plasmodia* within infected RBCs [17], was observed (Fig 2, bright field, and merge). The dark Hz pigment was visible inside the parasite-infected nucleated erythroid cells as well as in infected reticulocytes that originated from the donor. Hz is an insoluble, crystallized digestion product of heme derived from the digestion of hemoglobin by malaria parasites, containing heme-derived β-hematin, which neutralizes the toxicity of free heme released after parasite invasion through a digestive process that involves the digestive vacuole structure [18]. On days 3 to 6 post-infection, no parasite-infected erythroid cells were observed, and cell mortality had substantially increased. Therefore, the experiment was stopped on day 6.

**Fig 1.**
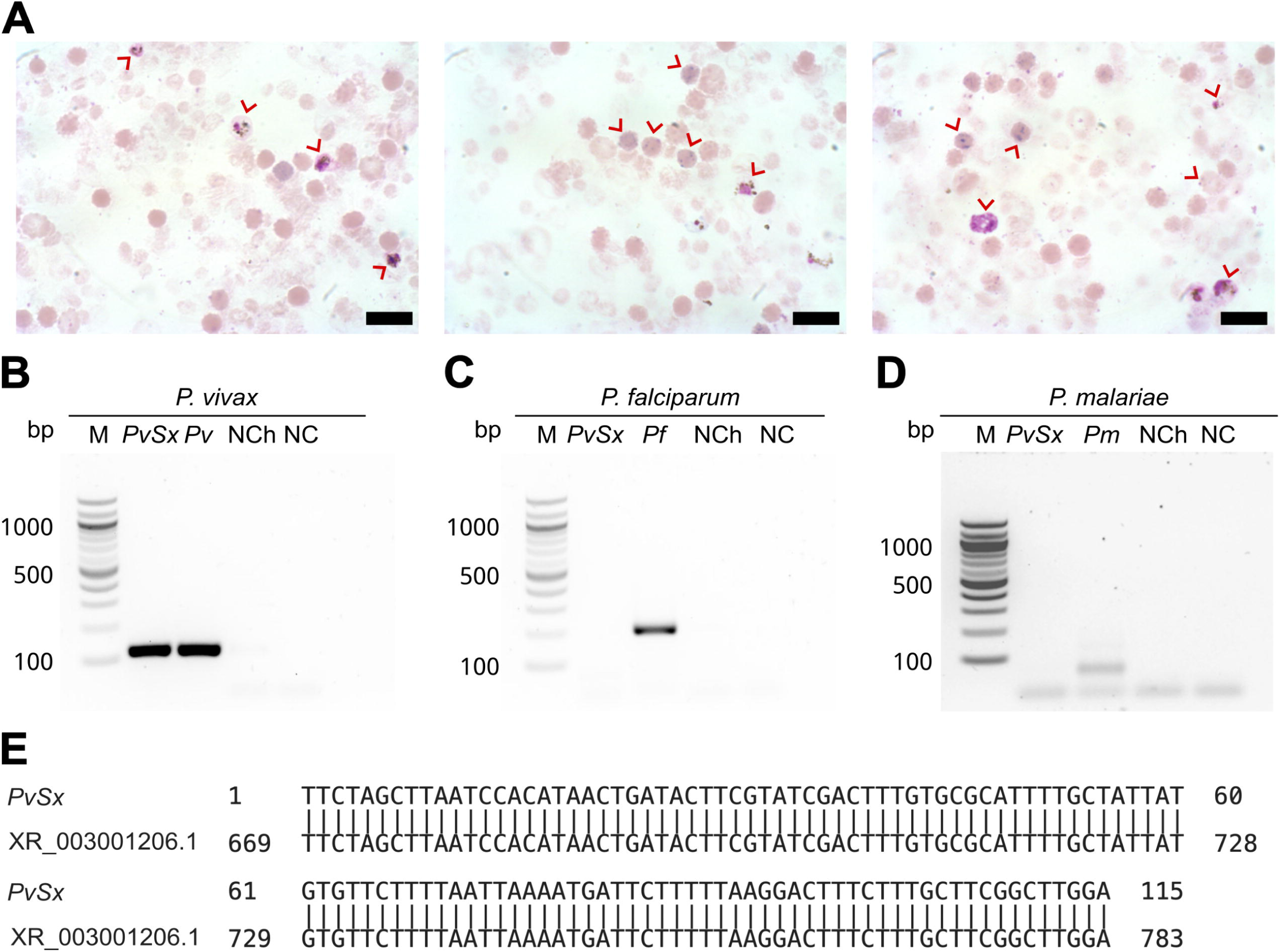
*P. vivax* mono-infected reticulocytes. (A) Three representative micrographs of *P. vivax*-infected enriched reticulocytes (red arrows) from a malaria patient. Scale bars are 3 µm. (B-D) Species-specific nested PCR for the small subunit 18S ribosomal RNA. Lane M, 100 bp marker; Lane *PvSx*, sample used in the infection of erythroid cells; Lanes *Pv*, *Pf*, and *Pm* belong to the positive controls for each species; lane NCh, negative control using genomic DNA from a healthy human; lane NC, negative control using water instead of genomic DNA. (B) *P. vivax* (∼120 bp amplicon). (C) *P. falciparum* (∼205 bp amplicon). (D) *P. malariae* (∼144 bp amplicon). (E) DNA sequence alignment of the positive *PvSx* amplicon with the corresponding gene segment of the *P. vivax* Salvador-1 reference strain (GenBank No. XR_003001206.1) [19], showing 100% identity.

**Fig 2.**
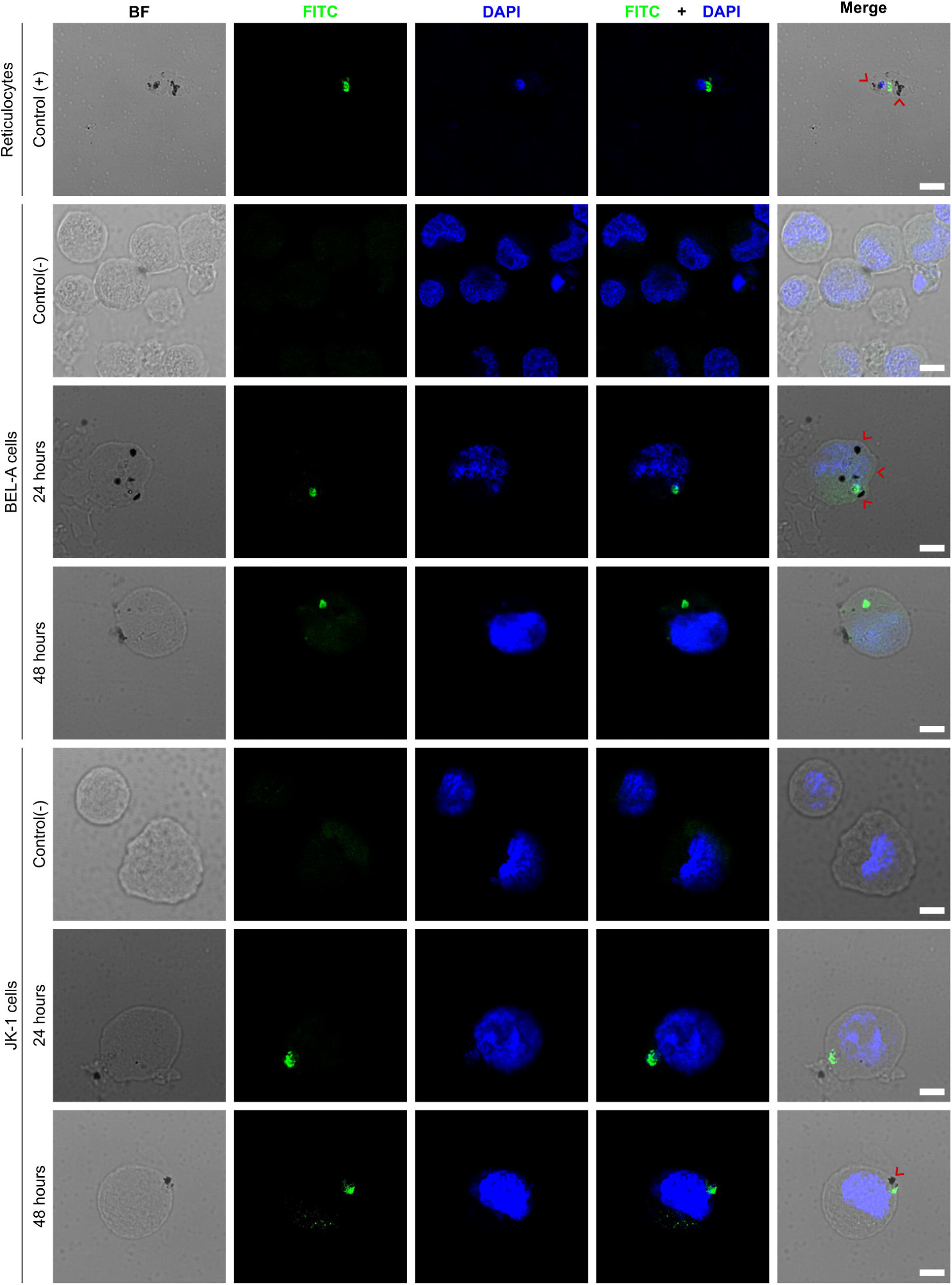
*P. vivax* invades BEL-A and JK-1 cells. The micrographs show *P. vivax*-infected reticulocytes as the positive Control (+); infected BEL-A and JK-1 cells at 24 and 48 h, and their non-infected negative Control (-); *P. vivax* lactate dehydrogenase (FITC green); DNA (DAPI blue); hemozoin crystals (black dots marked with red arrows); BF (bright field); Scale bars are 5 µm.

### Overlapping membrane proteomes reveal potential *P. vivax* invasion receptors

Because *P. vivax* was able to invade the erythroid BEL-A and JK-1 cells, their membranes must contain the same essential receptor molecules as reticulocytes that enable parasite invasion. Furthermore, the membranes of mature erythrocytes are expected to lack these receptors or to express them only at insufficient abundances. Consequently, a quantitative comparison of the membrane proteome of these cells with those of human reticulocytes and erythrocytes identified potential receptors for *P. vivax* merozoite ligands that are most likely responsible for its reticulocyte-restricted invasion. Stringent isolation procedures were necessary to obtain membrane proteins of pure reticulocytes. The isolated reticulocytes (CD71^+^, CD45^-^) used in this proteomic comparison had a purity of 98.4% (S2 Fig). Expression of CD71 is diminished during maturation into fully functional erythrocytes [20]. Simultaneous determination of CD45 negativity was necessary, as CD45^+^ leukocytes also express CD71, to ensure purity of the isolated reticulocytes.

The BEL-A and JK-1 cells used in the membrane proteomic comparisons were harvested from *in vitro* cultures and exhibited distinct nucleated erythroid maturation stages, including proerythroblasts, basophilic erythroblasts, polychromatic erythroblasts, and orthochromatic erythroblasts (S3 Fig), with slight dominance of the basophilic and polychromatic stages.

In total, 2,100 proteins were identified in BEL-A cells and 2,178 in JK-1 cells. The number of proteins was lower in reticulocytes (1,234) and in erythrocytes (1,347). After filtering this data for membrane proteins (see S1 Text & S4 Fig), 1,530 and 1,595 such proteins were obtained from BEL-A and JK-1 cell ghosts, respectively, while 846 and 974 proteins were identified for reticulocyte and erythrocyte ghosts.

Changes in membrane protein abundance were assessed by comparing erythroid cell lines and reticulocytes to mature erythrocytes. The protein abundancies of reticulocytes clustered better with those of BEL-A and JK-1 cells than with those of erythrocytes (Fig 3A). It was found that compared to erythrocytes 256 proteins were more abundant in reticulocytes, 1,179 in JK-1, and 1,554 in BEL-A. Of these, 144 identical proteins were increased in reticulocytes, JK-1, and BEL-A. However, reticulocytes and JK-1 cells share 18 proteins that are less abundant in erythrocytes, while BEL-A and reticulocytes have 40 proteins in common, that are less abundant in erythrocytes. Only 54 membrane proteins with higher abundance than in erythrocytes were identified exclusively in reticulocytes, 237 in JK-1, and 590 in BEL-A cells (Fig 3B).

**Fig 3.**
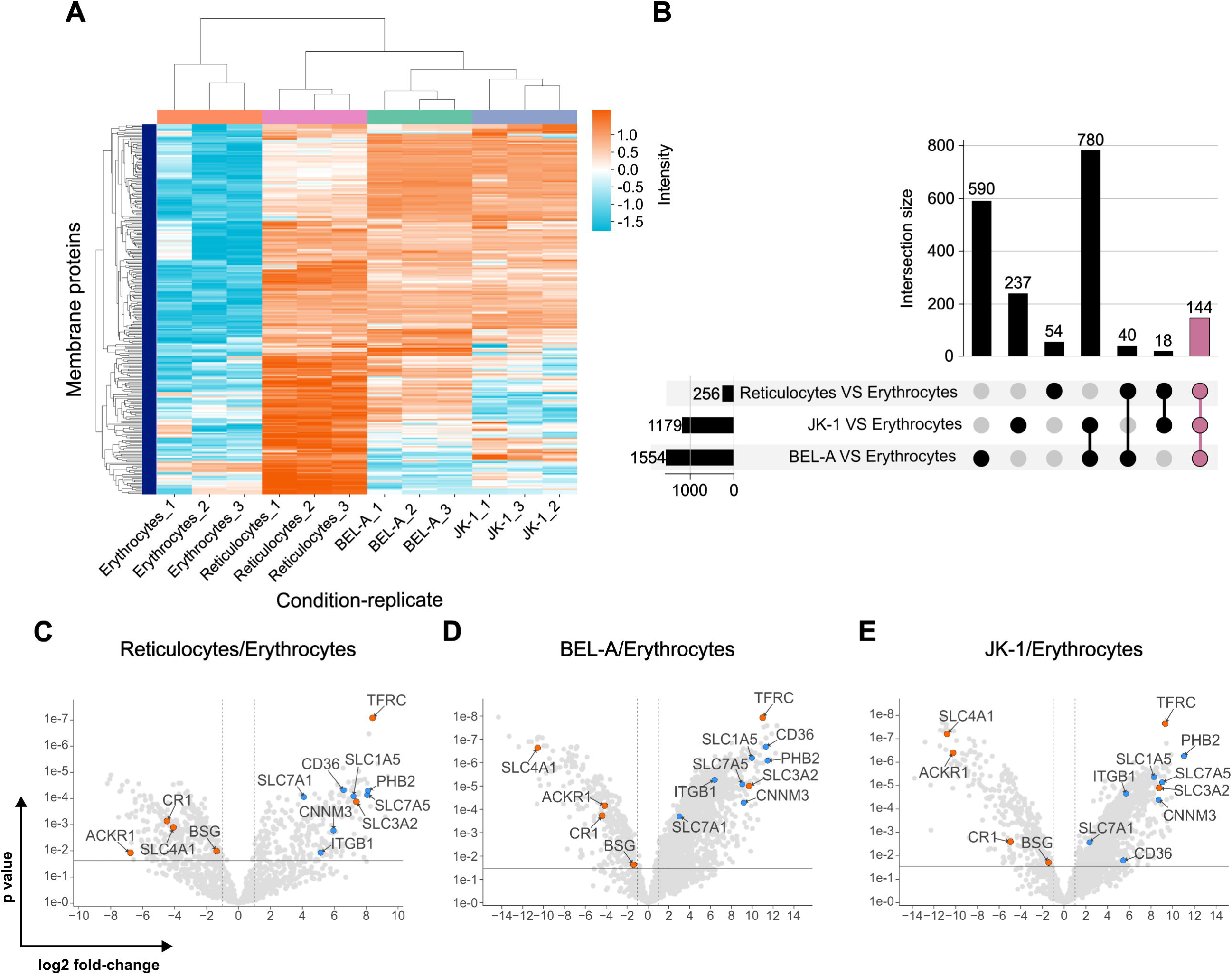
Plasma membrane proteomes of reticulocytes resemble those of erythroid cell lines. (A) Clustering of plasma membrane proteins in JK-1, BEL-A, reticulocytes, and erythrocytes, measured in triplicate. Log2 intensities. (B) Abundance of intersecting membrane proteins in reticulocytes, JK-1, and BEL-A cells compared to erythrocytes, represented in an UpSet plot [21]. 144 proteins share higher abundance among cell lines and reticulocytes but are reduced in erythrocytes (mauve bar). (C-E) Putative receptors (blue) and characterized receptors (orange) for *P. vivax* merozoite invasion. Gene names are displayed instead of protein names for simplicity. The x-axis represents the log_2_ fold change, and the y-axis shows the P-value, indicating statistical significance.

When comparing the membrane protein abundance in reticulocytes, cell lines, and erythrocytes, known *P. vivax* receptors such as TfR1 (CD71), CD98hc, ACKR1/DARC, BSG, CR1, and band 3 (SLC4A1) were identified. TfR1 and CD98hc, which are lost during reticulocyte maturation to erythrocytes, were significantly more abundant in reticulocytes and cell lines. In contrast, the other receptors showed higher levels in erythrocytes (Fig 3C-E).

*In silico* topological analysis of membrane proteins enriched in reticulocytes and erythroid cell lines identified several candidates — CD98lc (SLC7A5), high-affinity cationic amino acid transporter 1 (CAT-1, SLC7A1), neutral amino acid transporter B^0^ (ATB(0), SLC1A5), CD36, Integrin β-1 (ITGB1), prohibitin-2 (PHB2), and the metal transporter CNNM3 — as possessing sizable extracellular regions that are potentially accessible for interaction with *P. vivax* merozoite ligands (S1 Table & S5 Fig). The significantly higher abundance of these proteins in reticulocytes, BEL-A, and JK-1 cells compared to erythrocytes (Fig 3C-E & S1 Table) highlights them as potential candidates for *P. vivax* merozoite protein receptors, which may explain the parasite’s exclusivity for reticulocyte invasion.

### Receptors for PvRBP1a_158-650_LTID identified via proximity labeling

To identify potential receptors of the PvRBP1a_158-650_LTID protein, the TurboID proximity labeling technique was used. This technique enables the biotinylation of proteins that come into close contact with the fused protein (within 10 nm), facilitating the identification of their interactions [22,23]. Therefore, the fusion protein PvRBP1a_158-650_LTID and the LTID control were obtained in soluble form, with PvRBP1a_158-650_LTID having a molecular weight of ∼94 kDa and LTID ∼36 kDa (Fig 4A). Both proteins exhibited enzymatic activity and self-biotinylation at 1, 2, and 3 µM (Fig 4B-C).

**Fig 4.**
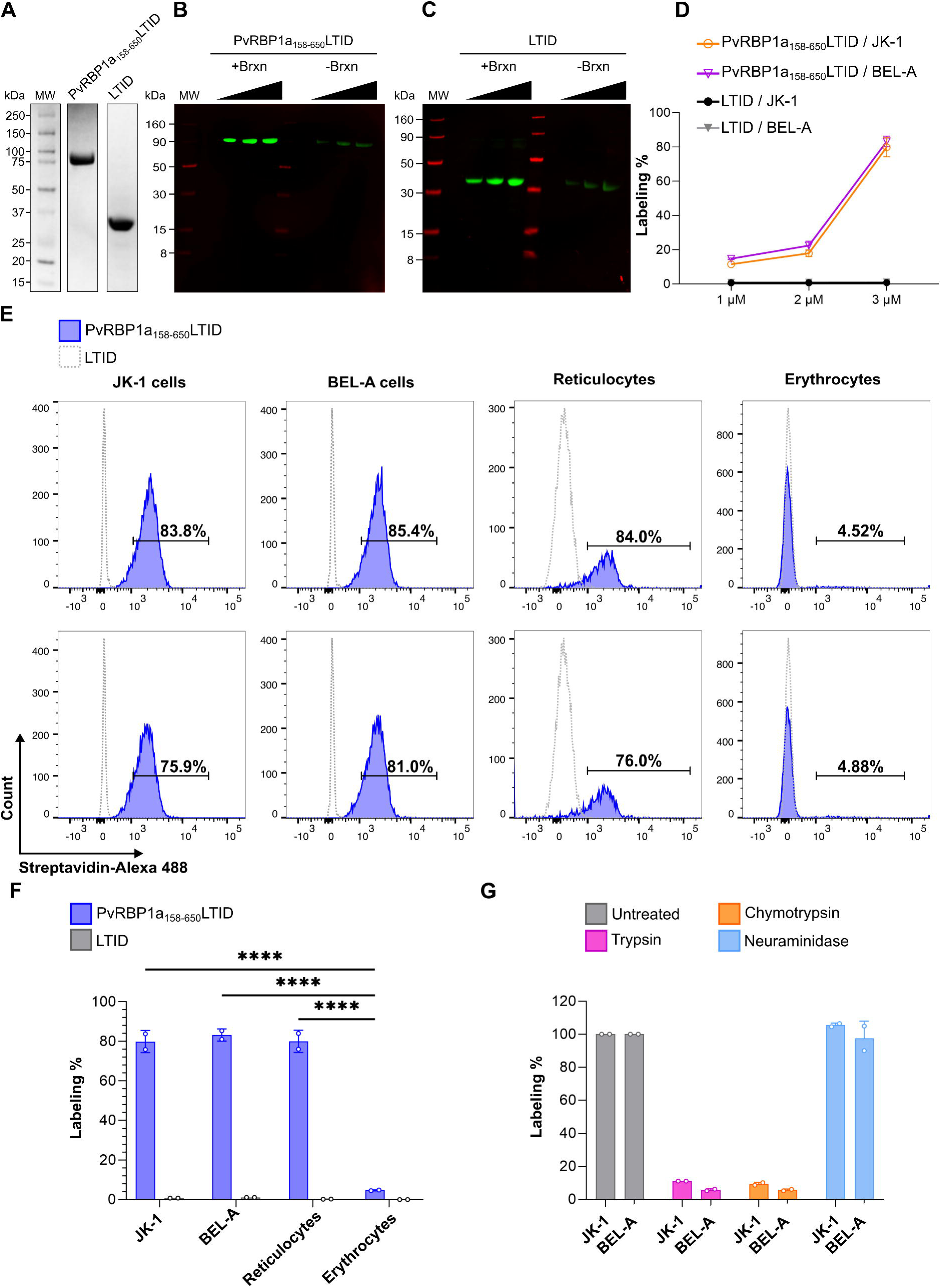
Proximity labeling with PvRBP1a_158-650_LTID in JK-1, BEL-A, reticulocytes, and erythrocytes. Recombinantly expressed soluble PvRBP1a_158-650_LTID and LTID by (A) SDS PAGE (B, C) auto-biotinylation assay with increasing concentrations of each protein at 1, 2, and 3 µM, detected with IRDye 800 Streptavidin (green bands), LICORbio molecular weight markers (MW, red bands); (D) biotin labeling of JK-1 (orange) and BEL-A (fuchsia) cells, in presence of PvRBP1a_158-650_LTID or LTID (negative control, JK-1 - black, and BEL-A - gray) at 1, 2, and 3 µM. (E-G) Flow cytometry (E) histograms of erythroid cells and human RBCs in presence of PvRBP1a_158-650_LTID (blue) or LTID (grey), both at 3 µM, demonstrating (F) significant degrees of cell surface labeling by PvRBP1a_158-650_LTID in erythroid cells and reticulocytes compared to erythrocytes (**** p ≤ 0.0001). (G) Receptors to PvRBP1a_158-650_ are sensitive to cell surface treatment with trypsin and chymotrypsin but resistant to neuraminidase (labeling % as normalized to untreated cells).

### PvRBP1a_158-650_LTID exhibits comparable binding to reticulocytes and erythroid cell lines via a proteinaceous receptor

Proximity biotinylation mediated by TurboID facilitated binding evaluation through the biotin-streptavidin interaction. The assays showed that PvRBP1a_158-650_LTID binding is concentration-dependent (Fig 4D). Since the highest percentage of biotin labeling on the cell surface was obtained at 3 µM, this concentration was selected to analyze PvRBP1a_158-650_LTID binding to enriched reticulocytes (87.5% purity) (S6 Fig), erythrocytes, and cell lines. The results showed that PvRBP1a_158-650_ had 80% biotinylation on the surface of reticulocytes, 4.7% on erythrocytes, and 79.85% and 83.2% on the surface of JK-1 and BEL-A cells, respectively. A statistically significant difference was found between the cell lines and reticulocytes compared to erythrocytes (P ≤ 0.0001). However, no significant difference was observed between the cell lines and reticulocytes (Fig 4E-F). These data suggest that erythroid cell lines exhibit PvRBP1a_158-650_ binding activity comparable to reticulocytes, indicating that they may express the receptor for this specific *P. vivax* ligand on their surface. Additionally, no labelling was detected with LTID, confirming the specificity of the PvRBP1a_158-650_ receptor interaction.

Cell surface labelling with PvRBP1a_158-650_LTID was sensitive to trypsin and chymotrypsin treatment but resistant to neuraminidase, which removes sialic acid from glycans that modify proteins in vertebrates (Fig 4G). Therefore, the cell surface receptor function for PvRBP1a_158-650_ is proteinaceous and not dependent on sialic acid-terminated glycans.

### Enrichment of PvRBP1a_158-650_LTID biotinylated cell surface proteins, identifies TfR1 and prohibitin-2 as the likely reticulocyte-restricting receptors

PvRBP1a_158-650_LTID biotinylated membrane proteins from erythroid cells were captured via streptavidin-affinity and subjected to proteomics analysis, revealing a total of 278 proteins, of which 12 were localized to the plasma membrane. However, the LTID control contained five of these proteins, leaving seven unique to enrichment after biotinylation with PvRBP1a_158-650_LTID. Four of these membrane proteins do not possess extracellular regions, while TfR1, prohibitin-2, and BSG do. Such extracellular regions should be required to facilitate an interaction with *P. vivax* merozoite ligands (Fig 5A). Interaction of PvRBP1a_158-650_ was successfully validated by targeted (PRM) LC-MS analysis for TfR1, BSG, and prohibitin-2 (Fig 5B). It clearly demonstrated that only PvRBP1a_158-650_LTID biotinylated these three proteins, while LTID did not.

**Fig 5.**
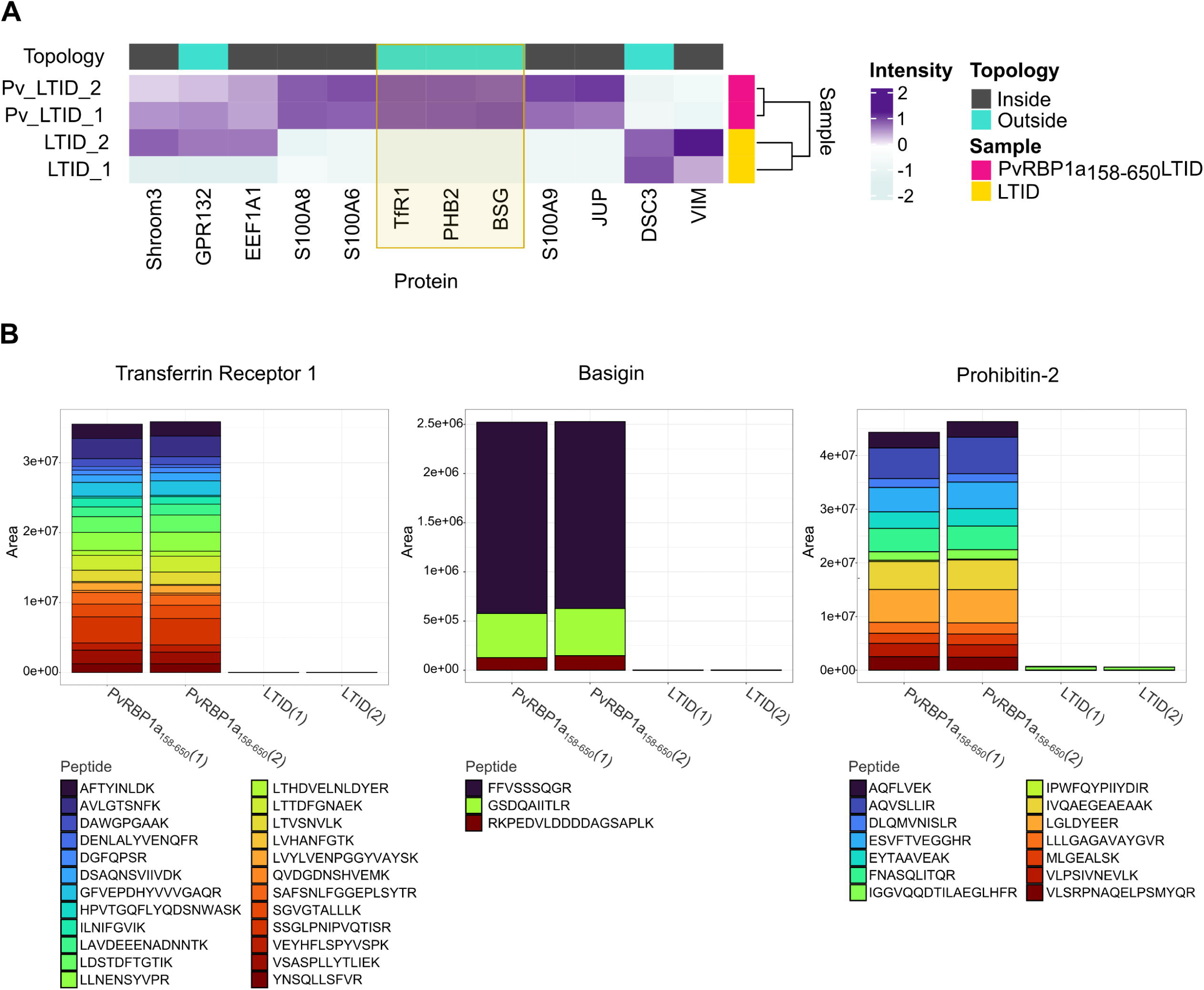
Proximity biotin-labeled TfR1 (CD71), Basigin, and Prohibitin-2, are identified as *P. vivax* receptor candidates after interaction of JK-1 cells with PvRBP1a_158-650_LTID and validated by targeted LC-MS. (A) Heatmap of labeled plasma membrane protein intensities in the PvRBP1a_158-650_LITD (fuchsia) and LTID (yellow) control samples, analyzed in duplicate by DDA LC-MS proteomics. Intensities are represented on a Z-score scale, where each value was transformed by the number of standard deviations (SD) from the mean. Topology categorizes proteins by their cellular localization: intracellular (Inside, dark gray) and those with extracellular domains (Outside, cyan). They include Shroom3, GPR132 – probable G-protein coupled receptor 132, EEF1A1 – Elongation factor 1-alpha 1, protein S100-A8, protein S100-A6, TfR1, PHB2 – prohibitin-2, BSG, protein S100-A9, JUP – junction plakoglobin, DSC3 – Desmocollin-3, and VIM – vimentin. (B) Validated interactions of transferrin receptor 1, basigin, and prohibitin-2 with PvRBP1a_158-650_LITD, but not with LITD, during the TurboID procedure, followed by PRM LC-MS. (1) (2) – duplicates. Stacked bars are the sum of the LC-MS ion chromatographic peak areas of the trypsin digested peptides (colored boxes) of each protein, indicating the contribution of each peptide to the individual protein abundance.

BSG is more abundant in erythrocytes than in reticulocytes and in the erythroid cell lines. In contrast, TfR1 and prohibitin-2 were significantly less abundant in erythrocytes (Fig 3C-E). In fact, TfR1 and prohibitin-2 were among the most abundant membrane proteins in reticulocytes and in the erythroid cell lines JK-1 and BEL-A. These findings suggest that PvRBP1a_158-650_ likely facilitate the recognition and invasion of reticulocytes through interaction with TfR1 and prohibitin-2.

### PvRBP1a_158-650_LTID interacts with high-affinity binding to TfR1, BSG, and prohibitin-2

The titration curves fit well to a single-site binding saturation model. In contrast, the negative control LTID displayed a nonspecific binding pattern “unstable”, corroborating the specificity of the interactions (Fig 6A). The Kd obtained from the ELISA titration indicated that PvRBP1a_158-650_LTID had high affinity for TfR1 (Kd: 1.15 nM), followed by BSG (Kd: 2.16 nM) and prohibitin-2 (Kd: 2.77 nM) (Fig 6B).

**Fig 6.**
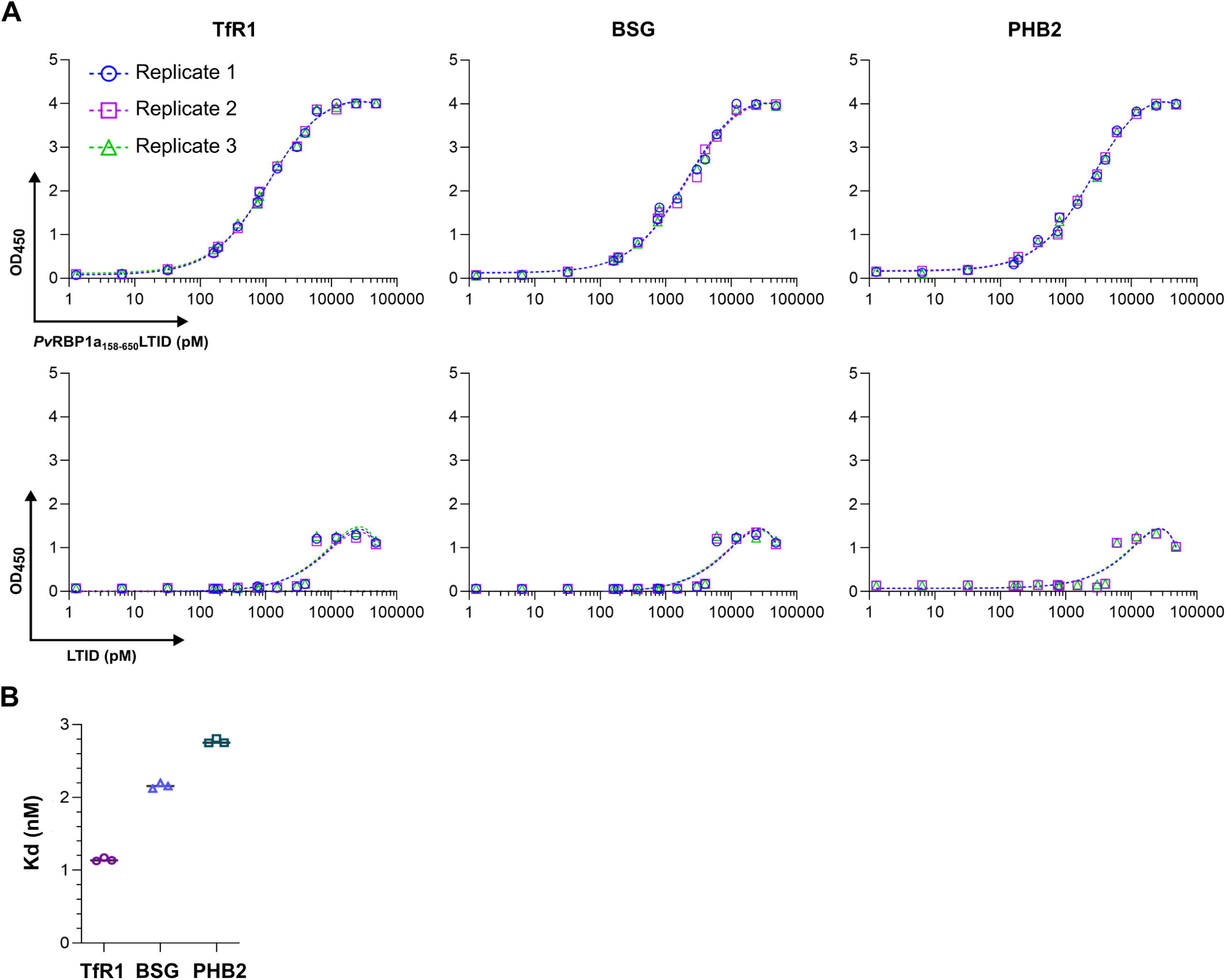
PvRBP1a_158-650_ binds to TfR1, BSG, and PHB2 at nanomolar affinities. (A) Titration ELISAs of protein-protein interactions between soluble ligand PvRBP1a_158-650_LTID, control LTID, and the immobilized receptor candidates TfR1, BSG, and PHB2. OD_450_ absorbance values represent the binding of a TurboID-specific HRP-labeled antibody for the biotin-free quantification of ligand and control in triplicate, fitted by a single-site binding model; (B) average dissociation constants (kd) summarized as determined from the fitted titration curves above.

## Discussion

This study demonstrates the utility of erythroid cell lines JK-1 and BEL-A as suitable surrogates for reticulocytes for studying the invasion process of the *P. vivax* malaria parasite. While these cell lines and their culture conditions did not support a continuous *P. vivax* culture *in vitro*, the formation of the Hz pigment and immuno detection of PvLDH strongly supported parasite invasion. In contrast to human reticulocytes, both erythroid cell lines are nucleated, which might permit them to initiate a cell-death program upon parasite invasion. Consistently, JK-1 cells were previously reported to support cell entry by both *P. vivax* and *P. falciparum* [24,25], while BEL-A cells have so far only been studied with *P. falciparum* [26]. This study is the first to report *P. vivax* invasion of BEL-A cells, confirming their susceptibility alongside JK-1 cells.

The quantitative membrane proteome comparison of reticulocytes and erythroid cell lines with erythrocytes revealed Prohibitin-2 (PHB2), TfR1 (CD71), the CD98 heavy chain (4F2hc, gene SLC3A2), the CD98 light chain (LAT-1, gene SLC7A5), ATB(0) (gene SLC1A5), CAT-1 (SLC7A1), CD36, Integrin β-1 (gene ITGB1), and Metal transporter CNNM3 to be of significantly increased abundance in those cell lines and reticulocytes. Whereas they are strongly decreased (practically absent) in fully matured erythrocytes. The increased abundance of TfR1 and CD98 in reticulocytes over erythrocytes is consistent with previous studies [2,20]. However, genetic manipulation [3] or antibody blockade [2] of these proteins only partially reduced *P. vivax* invasion, suggesting the involvement of additional receptors. The heavy chain of CD98 (SLC3A2) was reported to be bound by *P. vivax* in immature red blood cells via PvRBP2a [2]. Therefore, other potential *P. vivax* receptor candidates with extracellular regions, namely Prohibitin-2, the CD98 light chain (LAT-1), ATB(0), CAT-1, CD36, Integrin β-1, and CNNM3 should be considered. These membrane proteins participate in various protein-protein interactions that facilitate the entry of microorganisms into host cells [27–39]. Interestingly, LAT-1, that together with its heavy chain 4F2hc forms the heteromeric CD98 [40,41], plays a role in hepatitis C virus entry [42], raising the question of whether *P. vivax* may also interact with LAT-1.

Proximity labeling of erythroid cells with PvRBP1a_158-650_LTID was largely consistent with previous observations, in which ∼50% of reticulocytes binding and ∼20% of erythrocytes bound to PvRBP1a_157-650_ [13]. Furthermore, 20 of reticulocytes and only 1% of erythrocytes bound to PvRBP1a_351-599_ [43], while 31.5% of reticulocytes were reported to bind to PvRBP1a_30-778_ [44]. Additionally, the trypsin and chymotrypsin sensitivity of these recombinant protein [13,43,44], as well as the PvRBP1a_157-653_ HAPBs [12] align with our results.

PvRBP1a_158-650_ was found to interact with Prohibitin-2, TfR1, and BSG. Prohibitin-2 and TfR1 are more abundant in reticulocyte membranes and cell lines compared to erythrocytes, while BSG is more abundant in erythrocytes. These findings suggest that PvRBP1a_158-650_ may facilitate reticulocyte recognition and invasion through interaction with Prohibitin-2 and TfR1. Additionally, interaction with BSG may contribute to binding activity to erythrocytes, but not their restricted invasion, consistent with previous studies on PvRBP1a binding [12,13,43,44].

This study demonstrated strong binding of PvRBP1a_158-650_ with the 89-760 domain of TfR1, contrasting with prior work that did not detect this interaction, possibly due to the crucial role of TfR1’s 89-120 region, not included in the previous protein construct [25]. TfR1 is a known receptor for PvRBP2b [3], as well as for various New World arenaviruses [45,46]. BSG, a known receptor for *P. falciparum* RH5 [47,48], a member of the PfRH family homologous to the PvRBP proteins of *P. vivax* [49,50], also serves as a receptor for *P. vivax* TRAg38 [51]. Similarly, prohibitin-2, facilitates viral entry into host cells, including enteroviruses, coronaviruses, HIV-1, and flaviviruses such as dengue [39,52–54].

This research demonstrated the binding versatile of PvRBP1a_158-650_, which interacts with three different membrane proteins. In biological systems, ligands often bind multiple receptors, as seen with *Plasmodium* interactions; PvTRAg38 binds both BSG and band 3 [51,55], and PfEMP1binds to several receptors [56–60].

This study demonstrates that the BEL-A and JK-1 cells are suitable models for studying *P. vivax* receptor-ligand interactions, providing viable alternatives to reticulocytes. The similarity in the abundance of potential receptor candidates between cell lines and reticulocytes, and their dissimilarity with erythrocytes, validates the use of JK-1 and BEL-A cell lines as surrogate models for the study of *P. vivax* merozoite ligand-receptor interactions, and suggests the existence of other potential *P. vivax* receptors. prohibitin-2 and TfR1 may contribute to a redundant reticulocyte-restricted invasion pathway because they exhibit high binding affinities to PvRBP1a_158-650_. These findings lay the foundation for the comprehensive study of all *P. vivax* invasion mechanisms and for the development of targeted therapies against malaria.

## Materials and methods

### Ethics statement

The study was conducted in accordance with the Declaration of Helsinki. Use of anonymized discarded blood from therapeutic phlebotomy was approved by the Institutional Review Board of City of Hope, Duarte, California, USA, as exempt category 4, under 45CFR46.104 (d). Blood samples from malaria patients were obtained under informed consent with the approval of the Bioethics central committee of the Universidad de Córdoba, Monteria, Colombia, and imported into the United States under CDC permit No.: 20210830-3188A0.

### Collection, processing, and enrichment of *P. vivax* parasites from blood samples

*P. vivax* infected blood samples were collected from malaria patients in Tierralta— Córdoba, Colombia, into 5-mL sodium citrate tubes. After transportation to Bogotá, RBCs were enriched by centrifugation, mixed with an equal volume of Glycerolyte 57, cryopreserved, shipped to the U.S. lab, thawed using the NaCl method [61], and resuspended in 3 mL of Iscove’s Modified Dulbecco’s Medium (IMDM). These RBCs were then enriched from 0.2% to 4.0% parasitemia by concentrating *P. vivax*-infected reticulocytes through a KCl-Percoll gradient [62]. Enrichment was evaluated by microscopy with Giemsa staining (Fig 1A).

### Identification of the *Plasmodium* species

Genomic DNA was extracted from infected cells, and nested PCR was performed using this DNA to identify the *Plasmodium* species. Genus- and species-specific primers targeting the parasite’s 18S ribosomal small subunit RNA were used as previously described [63] (see S1 Text).

### *P. vivax* entry into BEL-A and JK-1 cells

The enriched *P. vivax* mono-infected RBC sample (∼40 µL) was divided into two equal aliquots, one for co-incubation with BEL-A cells and the other with JK-1 cells. Each cell line (1.5×10^5^ cells) was cultured in 500 µL of medium, for BEL-A in StemSpan serum-free expansion medium (SFEM, StemCell Technologies), containing 25% human serum (Type AB, Sigma-Aldrich), 50 ng/mL stem cell factor (SCF), 3 U/mL erythropoietin (EPO), 1 μM dexamethasone, 1 μg/mL doxycycline, 1 μg/mL chemically defined lipid concentrate (CDLC), and 100 µM hypoxanthine; and JK-1 in IMDM supplemented with GlutaMAX, with the same components, except for SCF, EPO, dexamethasone, and doxycycline. Both cultures were incubated at 37°C with 5% CO_2_ and 5% O_2_. Fresh medium and 2×10^5^ erythroid cells were added every two days, and cultures were evaluated by immunofluorescence assay (IFA) every 24 hours for 6 days (see S1 Text).

### Quantitative comparison of membrane proteomes by (DIA) - LC-MS/MS

Reticulocytes, erythrocytes, JK-1 and BEL-A cells were collected, cytoplasmic content was removed by osmolytic lysis, and membrane proteins of the remaining ghosts were extracted. From each sample, 23 μg of proteins were processed for proteomics using S-Trap columns (ProtiFi) according to the manufacturer instructions [64,65]. The resulting trypsin/LysC-digested peptides were analyzed by LC-MS/MS in data-independent acquisition (DIA) mode, as detailed in the S1 Text.

To enrich the plasma membrane proteins from the data set, proteins were filtered based on at least one of the following annotations from the UniProt subcellular localization database: “Cell membrane”, “Apical cell membrane”, “Basolateral cell membrane”, “Peripheral membrane protein” and “Plasma membrane”, and the Gene Ontology (GO) annotation term: “plasma membrane”. The topology of the membrane proteins abundant in reticulocytes compared to erythrocytes was evaluated using several predictors. Protein sequences were analyzed with Protter for overall visualization of proteoforms [66], TMHMM 2.0 for transmembrane region prediction [67], SignalP 6.0 for signal peptide identification [68], and PredGPI for GPI anchor site prediction [69].

### PvRBP1A_158-650_ proximity labeling for the identification of likely receptor candidates

#### Cloning, expression, purification, and activity of TurboID fusion proteins

The DNA sequence encoding PvRBP1a_158-650_ (GenBank AAS85749.1) was derived from the *P. vivax* Salvador I reference strain (txid126793). This sequence was fused to an acidic linker (L), GDEVDEDEG, to improve solubility, and the TurboID protein (TID) [22], followed by a C-terminal 6xHis tag for purification, resulting in the PvRBP1a_158-650_LTID fusion protein. An equivalent gene encoding L with TurboID alone (LTID) was designed as a negative control. Both gene constructs were obtained as customized synthetic genes, optimized for expression in *E. coli,* and cloned into a pET-28a(+) expression vector between its *Nco*I and *Xho*I sites. The constructs were expressed in soluble form in *E. coli* BL21 cells and purified by affinity chromatography, as detailed in the S1 Text. Protein purity and expression were verified by polyacrylamide gel electrophoresis. The biotinylation activity of both recombinant TurboID fusion proteins was confirmed by evaluating their autobiotinylation activity through incubation in the presence or absence of biotinylation reaction buffer Brxn (20 mM Tris-HCl, 500 µM biotin, 2.5 mM ATP, pH 7.5) at 37°C for 15 minutes, quenching on ice and Western blot analysis with streptavidin-IRDye 800CW conjugate (1:1,000) and an Odyssey DLx imaging system (LICORbio).

#### PvRBP1a_158-650_LTID proximity labeling

To evaluate PvRBP1a_158-650_LTID’s interaction with JK-1, BEL-A, reticulocytes, and erythrocytes, proximity labeling assays were performed in duplicate, and repeated up to three times. Cells were washed twice with PBS supplemented with 2% human serum (HS 2%) and incubated with either PvRBP1a_158-650_LTID or LTID (negative control) for 3 hours at room temperature with constant mild agitation at 10 rpm. Following incubation, cells were washed three times with HS 2% to remove unbound proteins, then incubated with Brxn for 15 minutes at 37°C. The reaction was stopped by cooling on ice for 5 minutes, and the samples were washed with cold HS 2% before labeling with Alexa Fluor 488-conjugated streptavidin (10 µg/mL, Invitrogen) for 1 hour at room temperature. Biotinylation was quantified by cytometry, acquiring 100,000 events per sample on a FACSAria Fusion (BD). Data were analyzed with FlowJo v10.8.1 [70], calculating the percentage of biotinylated cells relative to total cells. LTID-treated and unlabeled cells served as negative controls.

#### The biochemical nature of PvRBP1a_158-650_ receptors

JK-1 and BEL-A cells were treated with trypsin (1 mg/mL, Sigma-Aldrich), chymotrypsin (1 mg/mL, Sigma-Aldrich), or neuraminidase (50 mU, Roche) for 1 hour. After enzymatic treatment, proteolytic enzymes were inactivated with soy trypsin inhibitor (0.5 mg/mL, Sigma-Gibco) [71]. Proximity labeling assays were then performed as described above, using PvRBP1a_158-650_LTID or LTID (3 µM).

#### Affinity enrichment and LC-MS identification of PvRBP1a_158-650_ proximity-labeled receptor candidates

JK-1 cells were incubated with either PvRBP1a_158-650_LTID or LTID (negative control), each at 3 µM for 3 hours. After incubation, cells were washed three times with HS 2%, then incubated with 100 µL of Brxn for 15 minutes at 37°C. The reaction was stopped as described above, and cells were resuspended in 500 µL of IP-MS lysis buffer (MS-compatible Magnetic IP kit, streptavidin, Pierce, Thermo Scientific), incubated on ice for 30 minutes with intermittent vortexing every 5 minutes. After centrifugation, the lysate’s supernatant was collected and combined with Streptavidin magnetic beads (50 µL, Thermo Scientific), incubated for 1 hour at 21°C, and then overnight at 4°C, to enrich biotinylated membrane proteins. The beads were washed, and biotinylated proteins were eluted sequentially with 100 µL of 50 mM biotin, 100 µL of elution buffer, and 100 µL of 5% SDS at 95°C.

The eluted proteins were reduced, alkylated, and processed for proteomics using S-Trap spin columns (ProtiFi) according to the manufacturer’s instructions [64,65]. The resulting trypsin/LysC digested peptides were analyzed by LC-MS in data-dependent acquisition mode. Data analysis was performed using FragPipe v22.0 [72]. Candidate receptor proteins for PvRBP1a_158-650_ were selected based on the presence of extracellular regions that are favorable for ligand interaction, evaluated using UniProt GO annotations and the TMHMM 2.0 predictor [67]. Receptor candidates in the PvRBP1a_158-650_LTID sample that were detected in both duplicates and of significantly higher abundance (≥ 2 fold) compared to the negative control (LTID) were also considered. Subsequently, a parallel reaction monitoring (PRM) method was applied to validate and quantify the peptides of interest (as detailed in the S1 Text).

### Binding affinities of *Pv*RBP1a_158-650_ to select receptor candidates by ELISA

The recombinant extracellular protein domains of receptor candidates TfR1 (Cys^89^-Phe^760^, SinoBiological), BSG (Met^1^-His^205^, SinoBiological), and full-length PHB2 (Origene) were used to evaluate the interaction between the PvRBP1a_158-650_ and its binding membrane proteins (S1 Fig). Maxisorp plates were coated in triplicates with 5 μg/mL of each protein for 2 hours at room temperature and blocked with SuperBlock Buffer (Thermo Scientific). Serial dilutions of PvRBP1a_158-650_LTID and LTID were prepared in blocking solution, ranging from 48,000 pM to 187.5 pM (1:2 dilution) and 4,000 pM to 1.28 pM (1:5 dilution), and incubated for 16 hours at 4°C. Bound PvRBP1a_158-650_LTID and LTID were detected with a TurboID-specific polyclonal rabbit antibody (anti-BirA mutated/TurboID, Agrisera, 1:10,000). After five washes with PBST, 100 µL of 3,3’,5,5’,-Tetramethylbenzidine (TMB) substrate was added, and the reaction was stopped with 50 µL of 1 M phosphoric acid. Absorbance was measured at 450 nm. Dissociation constants (Kd) were determined using GraphPad Prism v10.3.1 with non-linear regression and a one-site binding saturation model.

## Supporting information

Graphical Abstract

Graphical Abstract

## Acknowledgments

The authors are grateful to NHS Blood and Transplant, especially Professor Jan Frayne, for providing the BEL-A cell line. “BEL-A cell lines were created by Professor Jan Frayne, Professor David Anstee and Dr Kongtana Trakarnsanga with funding from the Wellcome Trust (grant numbers 087430/Z/08 and 102610), NHS Blood and Transplant and Department of Health (England)”. Furthermore, they wish to extend their gratitude to the “Grupo de Investigaciones Microbiológicas y Biomédicas de Córdoba – GIMBIC” from the Universidad de Córdoba, Colombia, for providing the P. vivax samples. This study was funded in parts by the Memorial Hermann Foundation in support of MK and JMF. The authors dedicate this study to Professor Manuel Elkin Patarroyo (1946-2025), in loving memory.

## Supporting information captions

**S1 Text. Supporting methods.**

**S1 Table. *P. vivax* merozoite candidate receptors**

**S1 Fig. Purity Analysis of Recombinant Proteins.**

Recombinant proteins BSG, TfR1, and PHB2were separated on a 4–12% polyacrylamide gel and stained with Coomassie blue. The observed bands correspond to the expected molecular weights for each protein: BSG (35 kDa), TfR1 (∼77.4 kDa), and PHB2 (∼35.7 kDa), confirming their purity. The Kaleidoscope molecular weight marker (Bio-Rad) is included in the left lane as a reference.

**S2 Fig. Reticulocyte enrichment.**

(A, B) Reticulocyte percentage in the initial whole blood sample. (C, D) Reticulocyte percentage after Percoll density (70%-62%) enrichment. (E) Purity of reticulocytes (CD71^+^ CD45^-^) after sorting collection. This high-purity reticulocyte sample was not evaluated with brilliant cresyl blue due to the small total volume of sample obtained (∼ 50 µL). (B, D) Blood smears stained with brilliant cresyl blue to identify reticulocytes (red arrows). Scale bars are 10 µm.

**S3 Fig. Differentiation of erythroid cells.**

(A-D) BEL-A cells. (E-H) JK-1 cells. (A, E) Proerythroblasts (20-25 µm) were identified by their large nucleus with fine chromatin, prominent nucleoli, and basophilic cytoplasm. These cells progressed to (B, F) Basophilic erythroblasts (16-18 µm), characterized by a slightly smaller nucleus with condensed chromatin and cytoplasm with pronounced basophilia due to increased ribosomes synthesizing Hb. Subsequently, cells differentiated into (C, G) polychromatophilic erythroblasts (12-15 µm), which exhibited a more condensed nucleus with clusters of heterochromatin. The final nucleated stage, (D, H) orthochromatic erythroblast, indicated by the red arrow in D (10-12 µm), was noted for its pyknotic nucleus and a cytoplasm rich in acidophilic Hb. At this stage, the nucleus is expelled to form reticulocytes (8-10 µm), which then mature into erythrocytes (7-8 µm) through the final step of erythroid differentiation.

**S4 Fig. Gene Ontology (GO) term analysis of plasma membrane proteins.** Scatterplots showing the GO terms: cellular component, molecular function, and biological process of enriched plasma membrane proteins. The bubble color indicates the count of the GO term, which was obtained from the DAVID functional annotation charts. The size of the bubbles indicates the frequency of the GO term in the Gene Ontology Annotation (GOA) database. The axes of the plot have no intrinsic meaning. REVIGO (version 1.8.1) was used to generate these scatterplots by reducing the dimensionality of a pairwise semantic similarity matrix of GO terms using Multidimensional Scaling (MDS). The resulting projection is not nonlinear. The guiding principle is that semantically similar GO terms should remain close together in the plot.

**S5 Fig. Topology of *P. vivax* merozoite receptor candidates.**

(A-J) Predicted locations of potential parasite receptors are illustrated for the signal peptide (yellow), extracellular domain (blue), intracellular domain (mauve), and membrane-embedded domain (red).

**S6 Fig. Reticulocyte enrichment process.**

(A) Initial percentage of reticulocytes in the whole blood sample (0.11%). (B) Percentage of reticulocytes after enrichment using a 70%-62% Percoll density gradient (10.5%). (C) Final percentage of reticulocytes (CD45^-^ and CD98^+^) following FACS sorting (87.5%).

