## Supplementary figures and images for "The reticulocyte restriction: invasion ligand RBP1a of *Plasmodium vivax* targets human TfR1, prohibitin-2, and basigin"

### Graphical Abstract

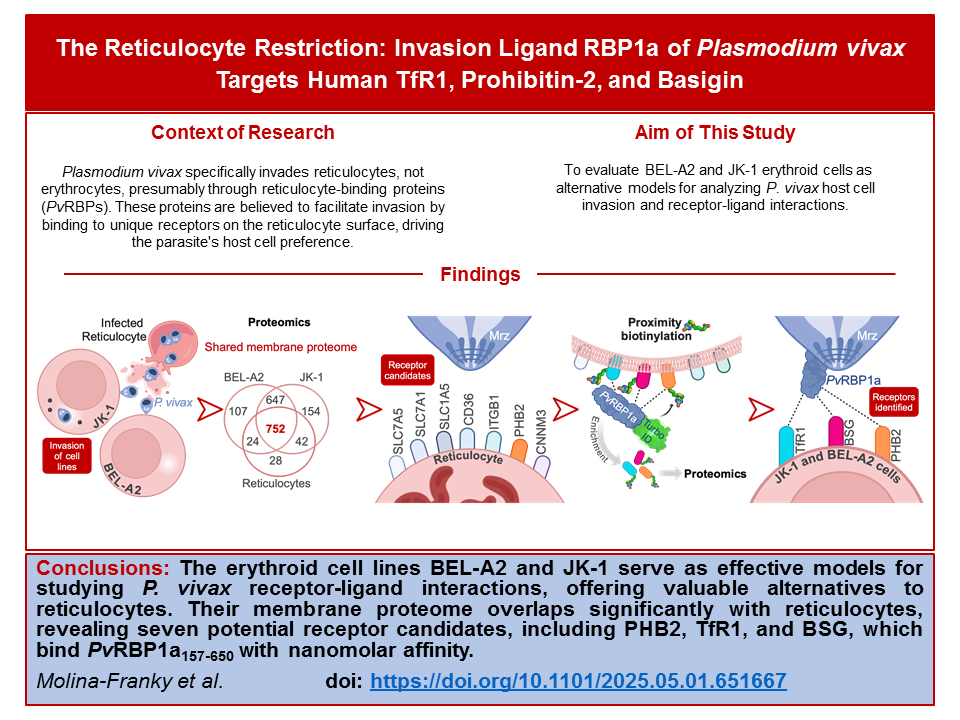

### S1 Fig

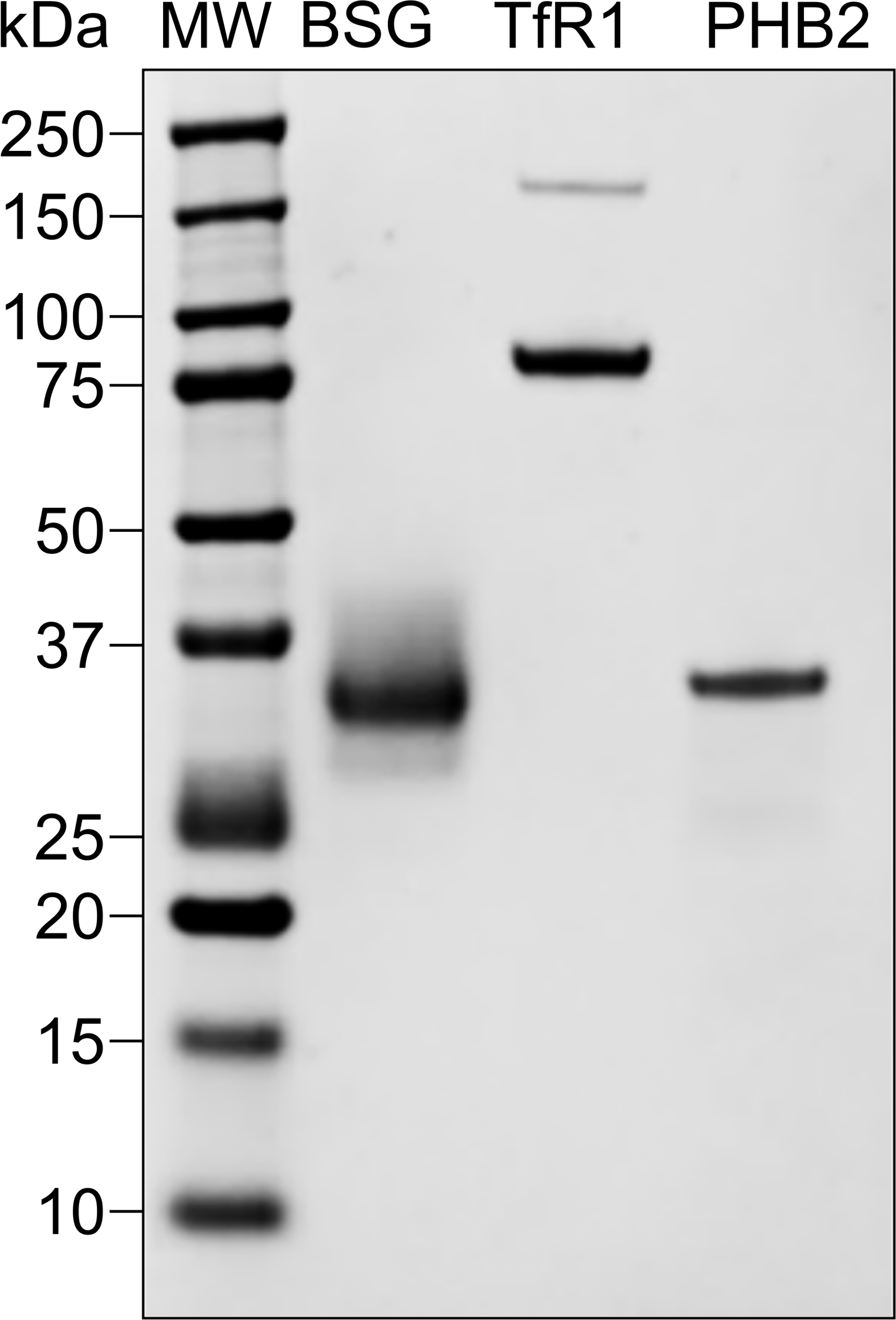

### S1 Fig

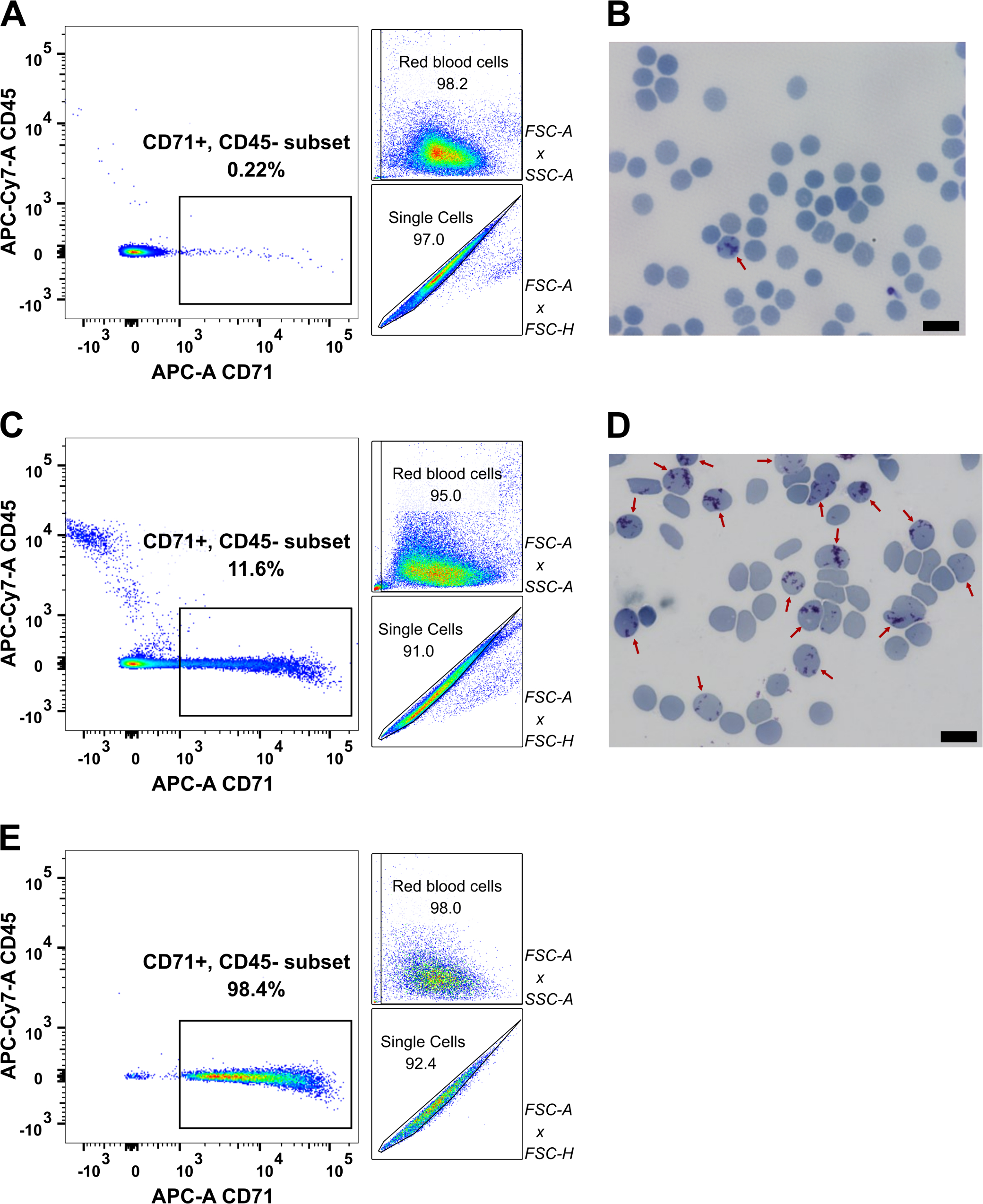

### S3 Fig

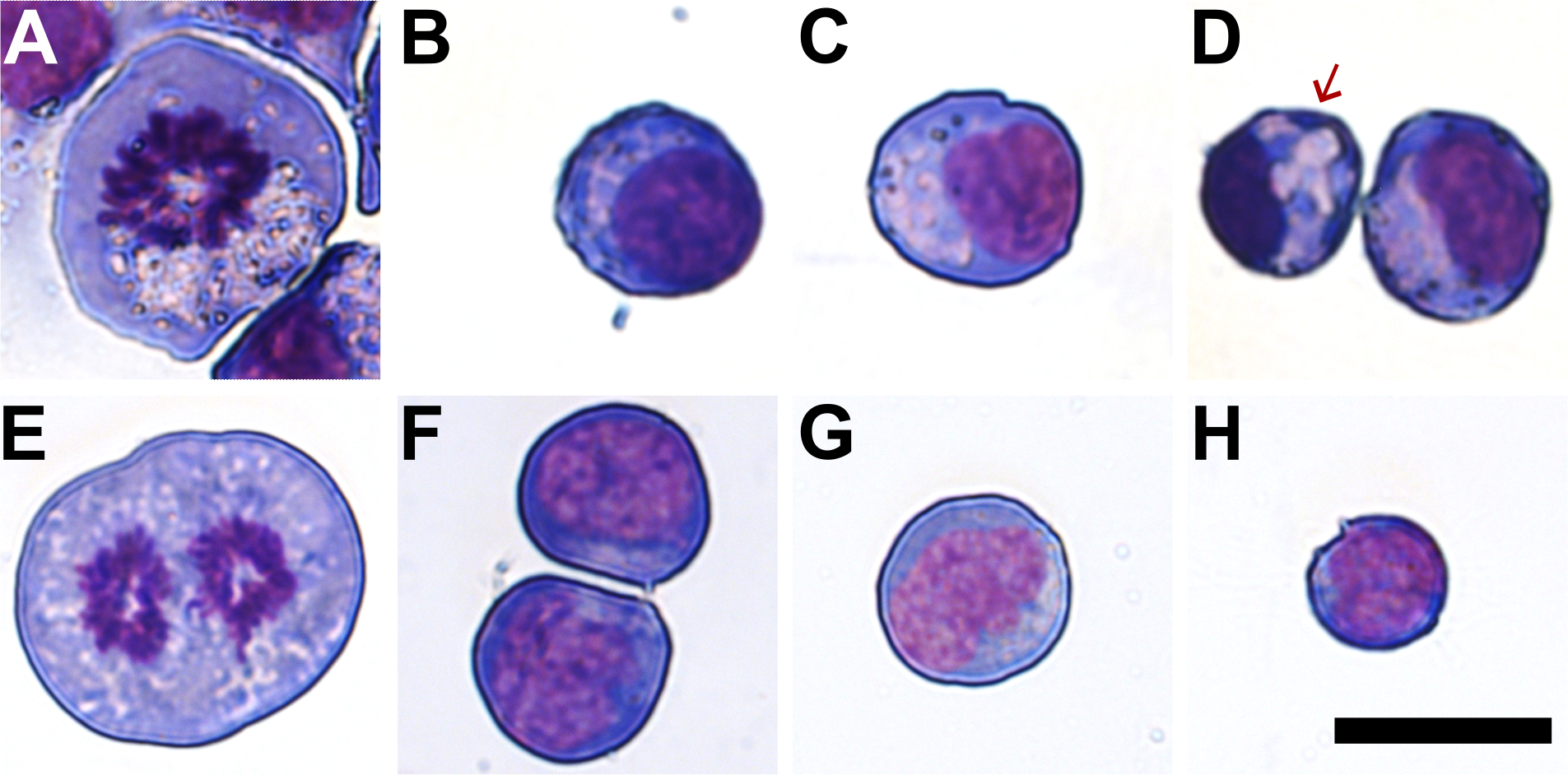

### S4 Fig

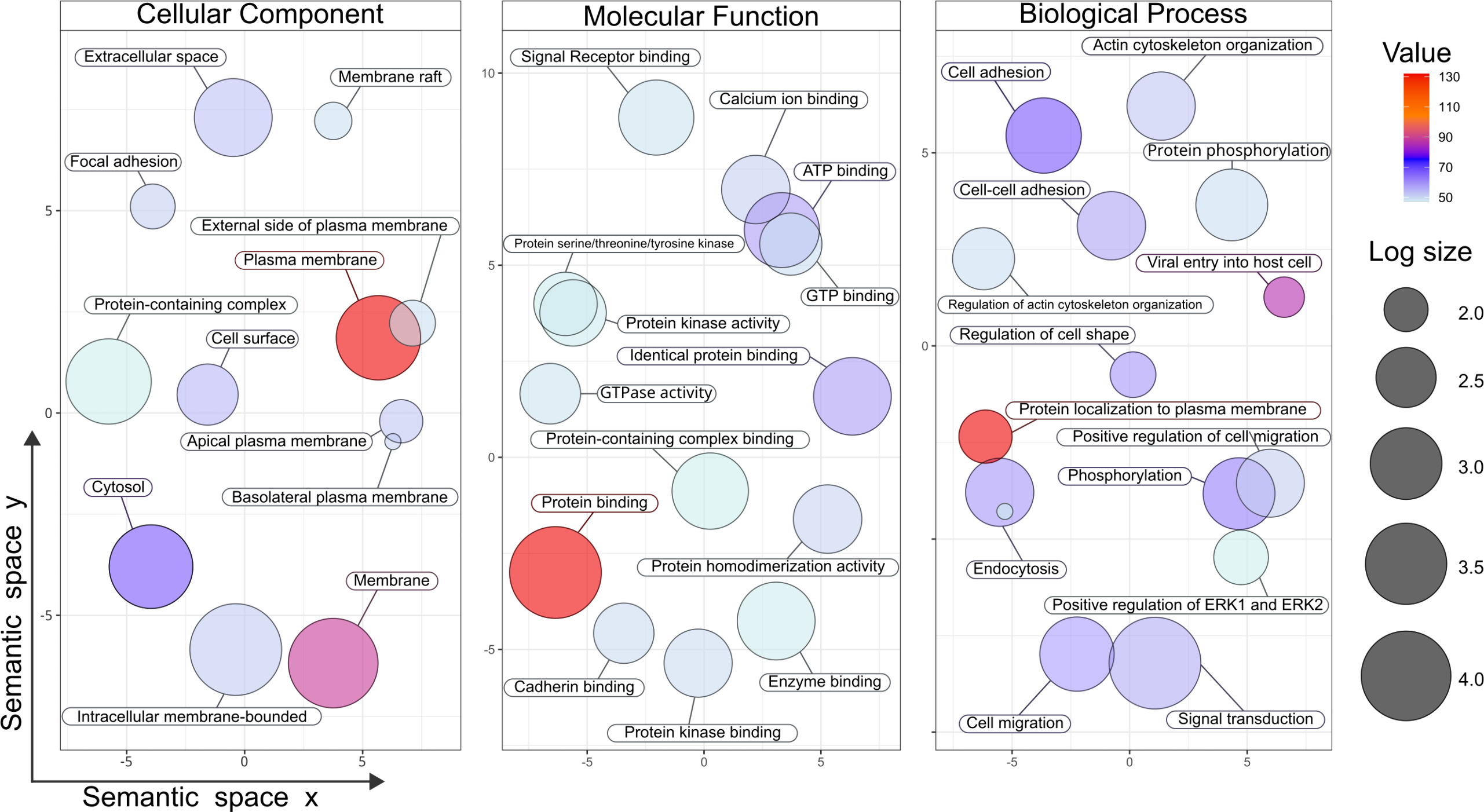

### S5 Fig

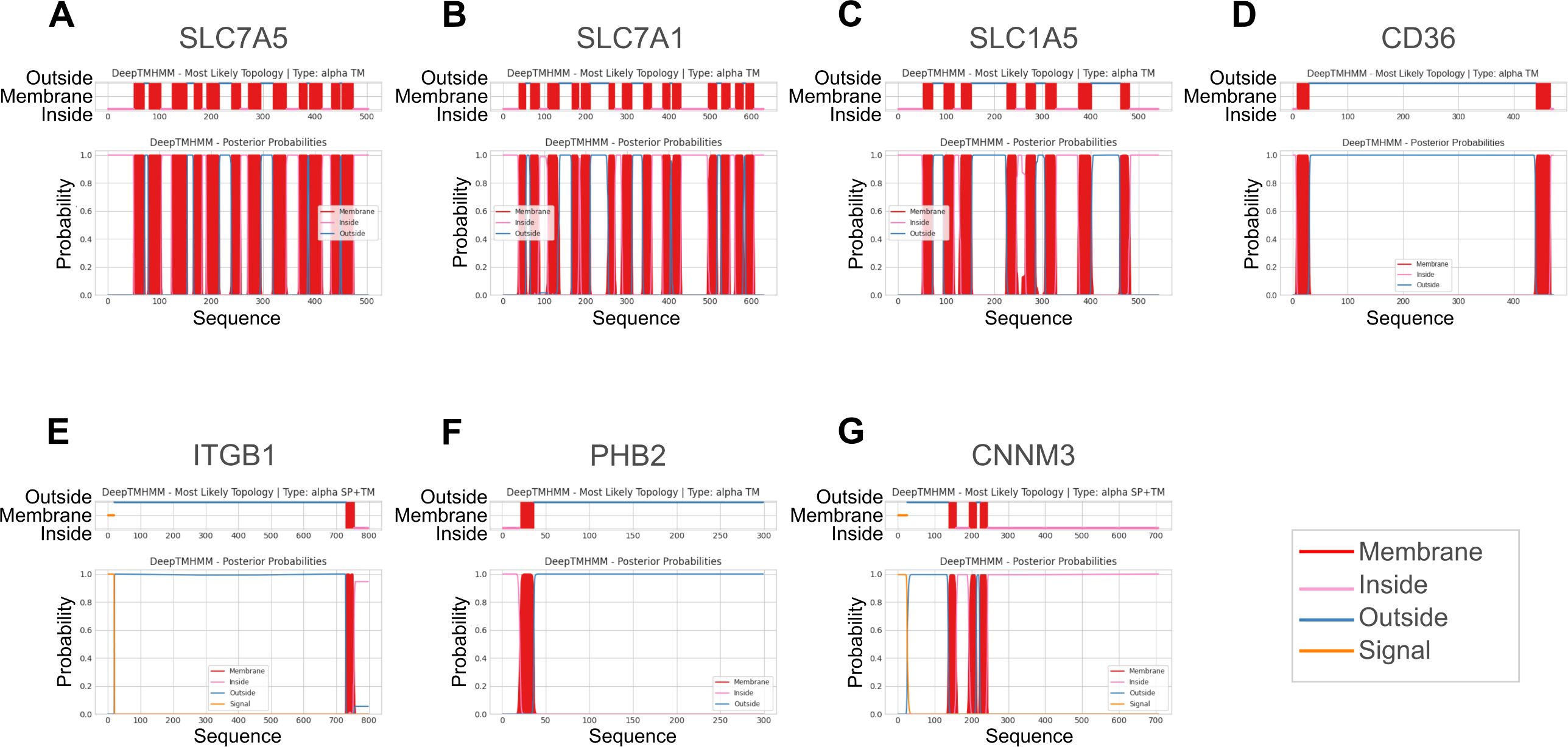

### S6 Fig

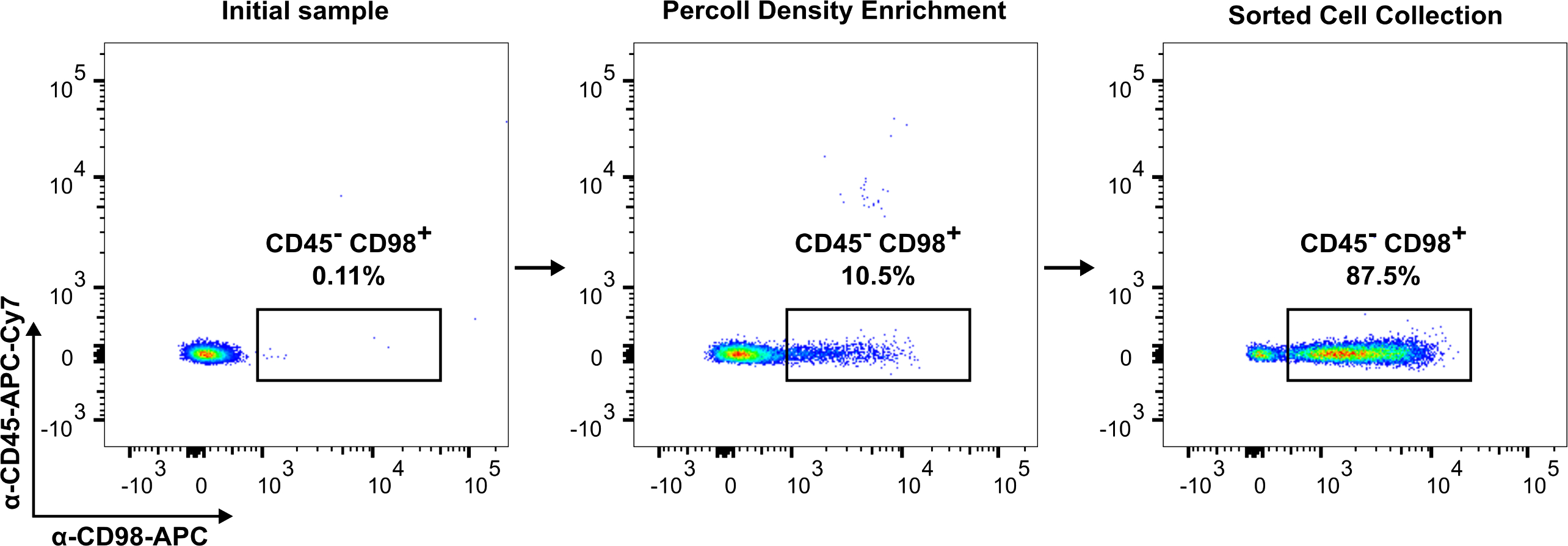
