## Supplementary material for "The reticulocyte restriction: invasion ligand RBP1a of *Plasmodium vivax* targets human TfR1, prohibitin-2, and basigin": S1 Text

**S1 Text. Supporting methods.**

**Detection of *Plasmodium* species by nested PCR**

1^st^ amplification: Each PCR reaction (25 μL total volume) contained primers rPLU 6 (TTAAAATTGTTGCAGTTAAACG) and rPLU 5 (CCTGTTGTTGCCTTAAACTTC) at 1µM each [1], - 5 µL of DNA template (100 ng), and GoTaq Green Master Mix (Promega, 1X). The conditions for the first amplification step were: 95°C for 5 minutes (1X); 94°C for 1 minute, 58°C for 2 minutes, 72°C for 2 minutes (30X); followed by 72°C for 5 minutes (1X).

2^nd^ amplification: Subsequently, the 1:35 diluted primary PCR product was used as the template in the secondary PCR, which was conducted with the species-specific primers:

rFAL 1 (TTAAACTGGTTTGGGAAAACCAAATATATT) and
rFAL 2 (ACACAATGAACTCAATCATGACTACCCGTC) for *P. falciparum*;

rVIV1 (CGCTTCTAGCTTAATCCACATAACTGATAC) and
rVIV2 (ACTTCCAAGCCGAAGCAAAGAAAGTCCTTA) for *P. vivax*;

rMAL 1 (ATAACATAGTTGTACGTTAAGAATAACCGC) and
rMAL 2 (AAAATTCCCATGCATAAAAAATTATACAAA) for *P. malariae*,
each at 1µM [1].

The PCR temperature profile was 94°C for 5 minutes (1X); 94°C for 30 seconds, 60°C for 1 minute, 72°C for 1 minute (35X); followed by 72°C for 5 minutes (1X). Genus-specific amplicons were separated by electrophoresis on a 2% agarose gel stained with SYBR Safe. The *P. vivax* amplicon was sequenced by Sanger and found to be identical with the *P. vivax* small subunit 18S ribosomal RNA (GenBank XR_003001206.1).

Positive control templates for *P. falciparum*, *P. vivax*, and *P. malariae*, and the negative control, healthy human genomic DNA, were from our validated collection.

**Immunofluorescence assay (IFA)**

BEL-A and JK-1 cells infected with *P. vivax* were fixed on immunofluorescence microscope slides with 4% paraformaldehyde in PBS. After washing and permeabilizing with 1% Triton X-100, cells were blocked with SuperBlock Buffer (Thermo Scientific) for 30 minutes. They were then incubated overnight at 4°C with mouse anti-*Pv*LDH antibody (1:100, The Native Antigen Company) in SuperBlock Buffer diluted 10-fold and supplemented with 0.05% Tween-20. After five washes with PBS, the slides were incubated with goat anti-mouse IgG –FITC antibody (1:200, Sigma-Aldrich) for 1 hour, washed again, and stained with 300 nM DAPI for 5 minutes. Cells were mounted in VECTASHIELD media, sealed, and imaged with a Zeiss LSM 880 confocal microscope. Images were processed utilizing ImageJ 1.53s [2].

**Collection of blood samples from non-infected humans for proteomics analysis**

500 mL of total peripheral blood were obtained from three patients undergoing therapeutic phlebotomy for high iron content. To separate the RBCs from the plasma and buffy coat, the blood was centrifuged at 1,900 × g for 5 minutes at 4°C, and the RBCs were washed three times in PBS. To limit proteolysis, the samples were kept at 4°C throughout every step.

To determine the percentage of reticulocytes, washed RBCs were diluted 1:10 with PBS and incubated with APC anti-Human CD71 (BD Pharmingen) and APC-Cy7 anti-Human CD45 (BD Pharmingen) antibodies (1:10 each) for 30 minutes at 4°C. The CD71^+^ CD45^-^ population was analyzed by flow cytometry using a FACSAria Fusion instrument (BD).

Reticulocytes were enriched by Percoll density gradient centrifugation, layering 6 mL of washed RBCs onto 6 mL of Percoll (3 mL 70% and 3 mL 62%). After centrifugation at 3,880 × g for 30 minutes at 4°C with brakes off, the reticulocyte-enriched fraction was collected, washed, and stained with anti-CD71 and anti-CD45 antibodies as described. The sample was then centrifugated at 150 × g, resuspended, and sorted on a FACS Aria Fusion (BD) to collect the CD71^+^ CD45^-^ population. Percoll-enriched and sorted samples were further evaluated by flow cytometry. The resulting purified reticulocytes were used to generate ghost reticulocytes (gRet).

For additional verification, both the initial and Percoll-enriched samples were stained with brilliant cresyl blue and analyzed under Bright-field microscopy at 63X.

Erythrocytes, depleted of reticulocytes, were collected after Percoll density gradient centrifugation. 5 mL of erythrocytes were washed with PBS, centrifuging at 1,900 × g for 5 minutes each time. Ghost erythrocytes (gEry) were subsequently prepared.

**JK-1 and BEL-A collection and morphological characterization**

JK-1 cells were cultured in IMDM GlutaMAX with 10% fetal bovine serum and 1X Penicillin-Streptomycin, while BEL-A cells were maintained in StemSpan SFEM medium with 50 ng/mL SCF, 3 U/mL EPO, 1μM dexamethasone, and 1μg/mL doxycycline. Both cell lines were cultured at 37°C with 5% CO_2_ at a cellular density of 70,000 – 150,000 cells/mL [3]. 1x10^7^ cells from each culture were harvested, washed with PBS, and used to prepare ghost cells (gJK-1 and gBEL-A).

For morphological characterization, 1x10^5^ cells of each line were seeded, fixed with methanol, and stained with Wright's eosin-methylene blue. Slides were analyzed by bright-field microscopy.

**Ghost cell preparation**

Reticulocytes, erythrocytes JK-1 and BEL-A cells were washed ~10 times and centrifuged at 17,000 × g for 15 minutes at 4°C with ghost lysis buffer (5 mM NaH₂PO₄, 10 mM NaCl, 0.5 mM EDTA and 1 mM PMSF, pH 8,0,) until white pellets of ghosts were obtained.

**Ghost cell protein extraction / membrane protein extraction**

gRet, gEry, gJK-1, and gBEL-A were incubated (1:1) with the protein extraction buffer (50 mM tetraethylammonium bromide (TEAB), 1X Halt protease inhibitor cocktail, 0.5 mM PMSF, 1% Triton X100 and 5% SDS) for 5 minutes at 95°C and centrifugated at 17,000 × g for 1 hour. Soluble proteins were recovered from the supernatants.

**Quantitative comparison of membrane proteomes by Data Independent Acquisition (DIA) LC-MS/MS**

Tryptic peptides were analyzed using an Exploris 480 orbitrap mass spectrometer with a 1200 Easy nanoLC system, using an Acclaim PepMap 100 pre-column (75µm × 2cm, nano Viper 2Pk C18, 3µm, 100 Å) and a PepMap RSLC C18 analytical reversed-phase column (75 µm × 25 cm, 2 µm, 100 Å Thermo, San Jose, CA, USA). Peptides were separated at a flowrate of 300 nL/min over a 195-minute gradient as follows: Buffer A consisted of 0.1% formic acid in water, and Buffer B was 80% acetonitrile with 0.1% formic acid. Gradient: 0 min at 2% B, 170 min to 33% B, 180 min to 100% B, and 195 min at 100% B. The full MS scan was acquired with an orbitrap resolution of 120K, and a range of 390-1100 m/z and an automatic gain control (AGC) target set to “standard”. Fragment ions were generated by high collision dissociation with 27% normalized collision energy and acquired by data-independently (DIA-MS/MS). The extraction window sizes were set to a width of m/z 8 and ranged from m/z 400 to m/z 1000. A second set of extraction windows was shifted by m/z 4 to achieve a staggered acquisition pattern as described by Pino et. al [4]. DIA spectra were acquired with a normalized AGC target of 1000% and a maximum ion injection time of 60 ms at 30K resolution.

**DIA LC-MS/MS Data Analysis**

Spectra in raw files were centroided, demultiplexed and files were converted to the mzML format using MSConvert 3.0.21101 [5]. Protein identification and quantitation was done with DIA-NN 1.8.2 [6] using the human UniProt reference proteome. Briefly, key parameters were: Enzymatic digestion with Trypsin, maximum of 2 missed cleavages, variable modifications for methionine oxidation and N-terminal acetylation, and fixed cysteine carbamidomethylation. The DIA-NN output files were processed and analyzed with R [7] and Mass Dynamics [8].

**Cloning, expression and purification of recombinant proteins**

Recombinant plasmids pET-28a(+)*Pv*RBP1a_158-650_LTID and pET-28a(+)LTID were obtained through gene synthesis services by Twist Bioscience (San Francisco, CA, USA), and transformed into *E. coli* BL21 competent cells (New England Biolabs). Each transformation involved incubating 100 ng of plasmid with cells on ice for 30 minutes, followed by heat shock at 42°C for 10 seconds and a final 5-minute incubation on ice. Cells were then recovered in 950 µL SOC medium, incubated at 37°C for 1 hour, and plated on Luria-Bertani (LB) agar containing kanamycin for selection. Plates were incubated for 16 hours at 37°C to confirm successful transformation.

Cells transformed with pET-28a(+)*Pv*RBP1a_158-650_LTID were cultured in 50 mL LB medium supplemented with 50 µg/mL kanamycin at 37°C and 250 rpm for 16 hours. The culture was diluted 1:10 (v/v) into 500 mL Terrific Broth (TB) with 50 µg/mL kanamycin and incubated under the same conditions. Upon reaching an OD600 of 0.7–0.8, the culture was cooled to 4°C for 30 minutes, and expression was induced with 0.5 mM isopropyl β-D-1-thiogalactopyranoside (IPTG) at 26°C and 220 rpm for 16 hours. Cells were harvested by centrifugation at 2,400 ×g for 20 minutes. For pET-28a(+)LTID-transformed cells, growth and expression conditions were identical except for IPTG induction at 1 mM, performed at 30°C for 4 hours. Harvesting was conducted as described above.

The expressed proteins were purified using affinity chromatography. The supernatants obtained from cell lysates were incubated for 16 hours at 4°C with 5 mL of nickel-NTA agarose resin (Thermo Scientific), pre-equilibrated with phosphate buffer at pH 8.0.

To minimize nonspecific binding and remove weakly bound proteins from the resin, the *Pv*RBP1a_158-650_LTID protein mixture was washed with 10 column volumes of phosphate buffer (pH 8.0) containing 10 mM imidazole and 0.1% Triton X-100, followed by an additional wash with 100 mL of the same buffer without detergent. Elution was performed using 10 mL of phosphate buffer (pH 8.0) with increasing concentrations of imidazole (40 mM, 250 mM, and 500 mM). Similarly, the LTID protein mixture underwent the same washing procedure, and elution was performed using phosphate buffer with increasing concentrations of imidazole (50 mM, 100 mM, 250 mM, and 500 mM).

Finally, *Pv*RBP1a_158-650_LTID and LTID proteins were buffer-exchanged to phosphate buffer at pH 7.5 using Amicon Ultra centrifugal filters with molecular weight cutoffs of 50 kDa and 30 kDa (Millipore), respectively. Protein quantification was conducted by densitometry, and the purified proteins were stored at -80°C.

**Reticulocyte purification for proximity labeling assays**

Peripheral blood (20 mL) was collected into EDTA tubes, and RBCs were separated from plasma and buffy coat by centrifugation at 1,900 × g for 5 minutes at 4°C. To determine the reticulocyte percentage, washed RBCs were diluted 1:10 in PBS and stained with anti-human CD98-APC (1:10, Miltenyi Biotec) and anti-human CD45-APC-Cy7 (1:10, BD Pharmingen) at 4°C in the dark for 30 minutes. CD98^+^ CD45^-^ cells were analyzed using a FACSAria Fusion (BD).

Reticulocytes were enriched by layering 6 mL of washed RBCs onto a Percoll gradient (3 mL of 70% and 3 mL of 62%) and centrifuging at 3,880 × g for 30 minutes at 4°C without applying the brake during deceleration to preserve cell integrity. The enriched reticulocyte fraction was collected, washed twice with PBS, and stained again with the same antibodies. After centrifugation at 150 × g for 5 minutes, the pellet was washed twice with PBS. CD98^+^ CD45^-^ reticulocytes were sorted using a FACSAria Fusion (BD).

**DDA LC-MS/MS Analysis of Proximity-labeled Biotinylated Membrane Proteins**

Digested peptides were analyzed using an Orbitrap Exploris 480 mass spectrometer (Thermo Fisher) equipped with an Easy nanoLC 1200 HPLC system. For separation, a pre-analytical Acclaim PepMap 100 column (75 µm × 2 cm, nano Viper C18, 3 µm, 100 Å) and an analytical PepMap RSLC C18 reversed-phase column (75 µm × 25 cm, 2 µm, 100 Å, Thermo, San Jose, CA, USA) were used. Peptides were separated at a flow rate of 300 nL/min over a 75-minute gradient. Buffer A consisted of 0.1% formic acid in water, while Buffer B was 80% acetonitrile with 0.1% formic acid. The elution gradient was as follows: 0 min at 2% Buffer B, 5 min at 2% Buffer B, 60 min at 38% Buffer B, 65 min at 100% Buffer B, and 75 min at 100% Buffer B. Spectra were acquired using a data-dependent acquisition (DDA) method, with full MS scans performed at a resolution of 60,000 in the range of 400–1,800 m/z. Ions with a charge state between 2 and 8 and a minimum intensity of 4E4 were selected for fragmentation, with an isolation window of 1 m/z. Fragmentation was carried out with a normalized collision energy of 28%, and the resulting spectra were recorded at a resolution of 30,000.

Data analysis was performed using FragPipe v22.0 [9], applying the default parameters for closed searches. The human proteome with isoforms from UniProt was used, and protein quantification was conducted using IonQuant. To enrich membrane proteins, the plasma membrane localization ontology from UniProt was utilized.

**Parallel Reaction Monitoring (PRM) Method**

To validate protein quantification results from DDA, samples were re-analyzed using PRM mass spectrometry. This targeted approach enhances the accuracy and specificity of protein quantification by selecting predefined ions for fragmentation, improving confidence in detecting and quantifying peptides of interest. The same column and gradient as in the DDA method were used to maintain consistency.

PRM analysis was performed using an inclusion list of unique peptides from candidate receptor proteins for *Pv*RBP1a_158-650_, including retention time, m/z ratio, and charge state from DDA experiments. Fragmentation was conducted within a 10-minute window around the average retention time from the DDA. MS/MS scans were acquired at a resolution of 30,000. Data analysis was performed with Skyline (64-bit) 24.1 [10], and results were summarized and visualized using R [7].
