## Supplementary material for "The reticulocyte restriction: invasion ligand RBP1a of *Plasmodium vivax* targets human TfR1, prohibitin-2, and basigin": S1 Table

**S1 Table. *P. vivax* merozoite candidate receptors.**

| **Gene** | **Protein name** | **Length**  **(aa)** | **TM Helix location** | **Extracellular regions** | **Signal peptide**  **location**  **(Sec/SPI probability)** | **GPI/**  **non GPI**  **anchor protein**  **Yes/No** | **Ret/Ery** | | **BEL-A/Ery** | | **JK-1/Ery** | |
| --- | --- | --- | --- | --- | --- | --- | --- | --- | --- | --- | --- | --- |
|  |  |  |  |  |  |  | **Log2 Fold change** | **Adjusted *P* value** | **Log2 Fold change** | **Adjusted *P* value** | **Log2 Fold change** | **Adjusted *P* value** |
| SLC7A5 | Large neutral amino acids transporter small subunit 1 | 507 | 55-75, 83-107, 128-157, 170-186, 195-219, 243-260, 275-299, 323-348, 373-389, 393-417, 435-452, 455-478 | 76-82, 158-169, 220-242, 300-322, 390-392, 453-454 | Non-SP  (0.0006) | No | 8.06 | 0.0019 | 9.06 | 7.98E^-05^ | 9.02 | 0.0001 |
| SLC7A1 | High affinity cationic amino acid transporter 1 | 629 | 38-55, 66-88, 108-136, 166-183, 188-210, 255-269, 287-311, 338-358, 385-403, 408-429, 495-516, 527-547, 560-579, 585-605 | 56-65, 137-165, 211-254, 312-337, 404-407, 517-526, 580-584 | Non-SP  (0.0001) | No | 4.09 | 0.0019 | 3.03 | 0.0007 | 2.33 | 0.0071 |
| SLC1A5 | Neutral amino acid transporter B(0) | 541 | 52-72, 95-116, 131-153, 226- 245, 265-285, 306-328, 374-401, 461-481 | 73-94, 154-225, 286-305, 402-460 | Non-SP  (0.0002) | No | 6.83 | 0.0023 | 9.26 | 4.34E^-05^ | 8.17 | 0.0003 |
| CD36 | Platelet glycoprotein 4 | 472 | 8-30, 440-466 | 31-439 | Non-SP  (0) | No | 6.57 | 0.0019 | 11.30 | 2.68E^-05^ | 4.10 | 0.009 |
| ITGB1 | Integrin beta | 798 | 729-754 | 21-728 | 1-20  (0.9992) | No | 5.13 | 0.0296 | 6.62 | 0.0005 | 5.71 | 0.0003 |
| PHB2 | Prohibitin-2 | 299 | 21-36 | 37-299 | Non-SP  (0) | No | 8.12 | 0.0019 | 6.09 | 6.36E^-05^ | 11.05 | 4.91E^-05^ |
| CNNM3 | Metal transporter CNNM3 | 707 | 138-158, 193-213, 223-243 | 26-137, 214-222 | 1-25  (0.999) | No | 5.95 | 0.0072 | 4.55 | 0.0004 | 8.71 | 0.0004 |
