## Supplementary material for "The reticulocyte restriction: invasion ligand RBP1a of *Plasmodium vivax* targets human TfR1, prohibitin-2, and basigin": Graphical Abstract

### Context of Research

*Plasmodium vivax* specifically invades reticulocytes, not erythrocytes, presumably through reticulocyte-binding proteins (*Pv*RBPs). These proteins are believed to facilitate invasion by binding to unique receptors on the reticulocyte surface, driving the parasite's host cell preference.

### Aim of This Study

To evaluate BEL-A2 and JK-1 erythroid cells as alternative models for analyzing *P. vivax* host cell invasion and receptor-ligand interactions.

### Findings

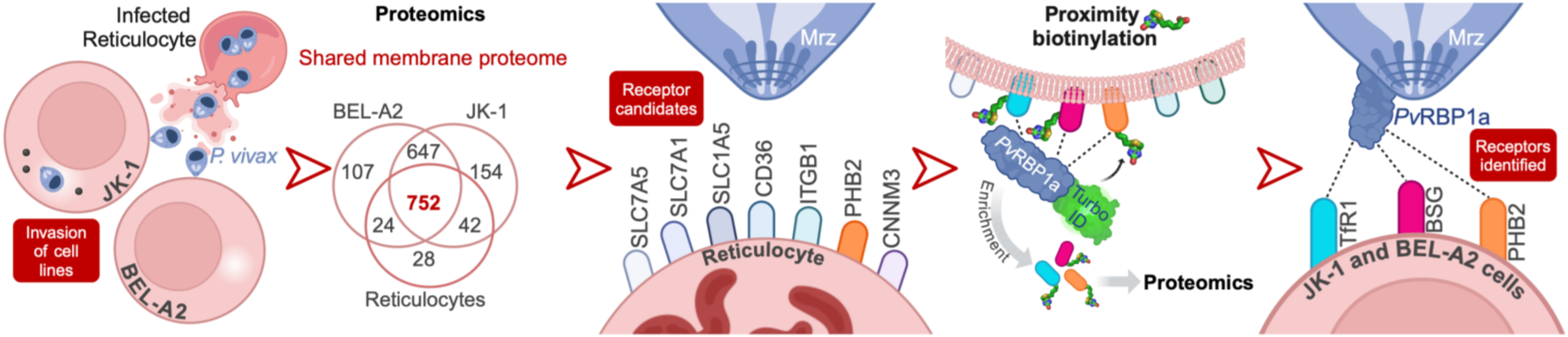


**Conclusions:** **The erythroid cell lines BEL-A2 and JK-1 serve as effective models for studying *P. vivax* receptor-ligand interactions, offering valuable alternatives to reticulocytes. Their membrane proteome overlaps significantly with reticulocytes, revealing seven potential receptor candidates, including PHB2, TfR1, and BSG, which bind *Pv*RBP1a$_{157\text{-}650}$ with nanomolar affinity.**

*Molina-Franky et al.*

doi: https://doi.org/10.1101/2025.05.01.651667
